# Structural and mutational analysis of the ribosome-arresting human XBP1u

**DOI:** 10.1101/588681

**Authors:** Vivekanandan Shanmuganathan, Nina Schiller, Anastasia Magoulopoulou, Jingdong Cheng, Katharina Braunger, Florian Cymer, Otto Berninghausen, Birgitta Beatrix, Kenji Kohno, Gunnar von Heijne, Roland Beckmann

## Abstract

XBP1u, a central component of the unfolded protein response (UPR), is a mammalian protein containing a functionally critical translational arrest peptide (AP). Here, we present a 3 Å cryo-EM structure of the stalled human XBP1u AP. It forms a unique turn in the upper part of the ribosomal exit tunnel and causes a subtle distortion of the peptidyl transferase center, explaining the temporary translational arrest induced by XBP1u. During ribosomal pausing the hydrophobic region 2 (HR2) of XBP1u is recognized by SRP, but fails to efficiently gate the Sec61 translocon. An exhaustive mutagenesis scan of the XBP1u AP revealed that only 10 out of 21 mutagenized positions in the XBP1u AP are optimal with respect to translational arrest activity. Thus, XBP1u has evolved to induce an intermediate level of translational arrest, allowing efficient targeting by SRP without activating the Sec61 channel and thereby serving its central function in the UPR.

## Introduction

Most secretory and membrane proteins traverse the endoplasmic reticulum (ER), where they are modified and folded before continuing their journey towards their respective destinations. The ER handles approximately one-third of the proteome and the flux of proteins entering the ER lumen varies widely, primarily depending on the demands of the specific cell type. The ER is also responsible for maintaining calcium homeostasis and is involved in lipid biosynthesis (Fagone and Jackowski, 2009; Görlach et al., 2006). A number of circumstances can alter the folding and modifying capacity of the ER, such as glucose deprivation, calcium imbalance, hypoxia, or viral infection, and thereby alter the demand on ER activity, as shown, e.g., in B cell differentiation (Grootjans et al., 2016). The central cellular response mechanism that alleviates ER stress and adjusts ER activity levels is the unfolded protein response (UPR) (Walter and Ron, 2011). In mammalian cells, this pathway is mainly mediated by three transmembrane sensors that are located in the ER membrane: inositol requiring enzyme 1 alpha (IRE1α), activating transcription factor 6 (ATF6), and pancreatic endoplasmic reticulum kinase (PERK) (Walter and Ron, 2011).

Of these three sensors, the evolutionarily most conserved is IRE1 (here, IRE1 denotes mammalian IRE1α and/or yeast Ire1); in lower eukaryotes such as yeast, it is the only known sensor mediating the UPR (Mori, 2009). IRE1 is a single-spanning membrane protein with three domains: a luminal unfolded protein-sensing domain and cytosolic bifunctional serine/threonine kinase and endo-ribonuclease domains. In unstressed cells, Hsp70 family chaperone BiP binds the luminal region of IRE1 and keeps IRE1 in an inactive monomeric state. Increasing levels of misfolded proteins during ER stress sequester BiP away from, leading to active dimer (Bertolotti et al., 2000; Okamura et al., 2000) and further highly activated by cluster formation (Aragón et al., 2009; Credle et al., 2005; Kimata et al., 2007; Korennykh et al., 2009; Li et al., 2010) In yeast, direct binding of unfolded proteins to the luminal core regions of IRE1-dimer or -oligomer is required for the activation (Gardner and Walter, 2011; Kimata et al., 2007), however, in mammals, direct binding model has been a matter of debate (Kohno, 2010). The cytosolic domain of activated IRE1α then splices the *XBP1u* (X-box binding protein-1 unspliced) mRNA on the ER membrane, producing *XBP1s* (spliced) mRNA, which codes for an active transcription factor. The splicing reaction involves removal of a 26-nt intron from *XBP1u* mRNA, which leads to a translational frame-shift and the replacement of C-terminal segment in XBP1u downstream of the splicing site (Calfon et al., 2002; Yoshida et al., 2001). Once translocated to the nucleus, the XBP1s transcription factor activates genes encoding ER-resident chaperones and folding enzymes, the components of ER associated protein degradation and the proteins that function in secretory pathway, which together increase ER size and activity (Figure 1) (Shaffer et al., 2004; Sriburi et al., 2004).

**Figure 1.**
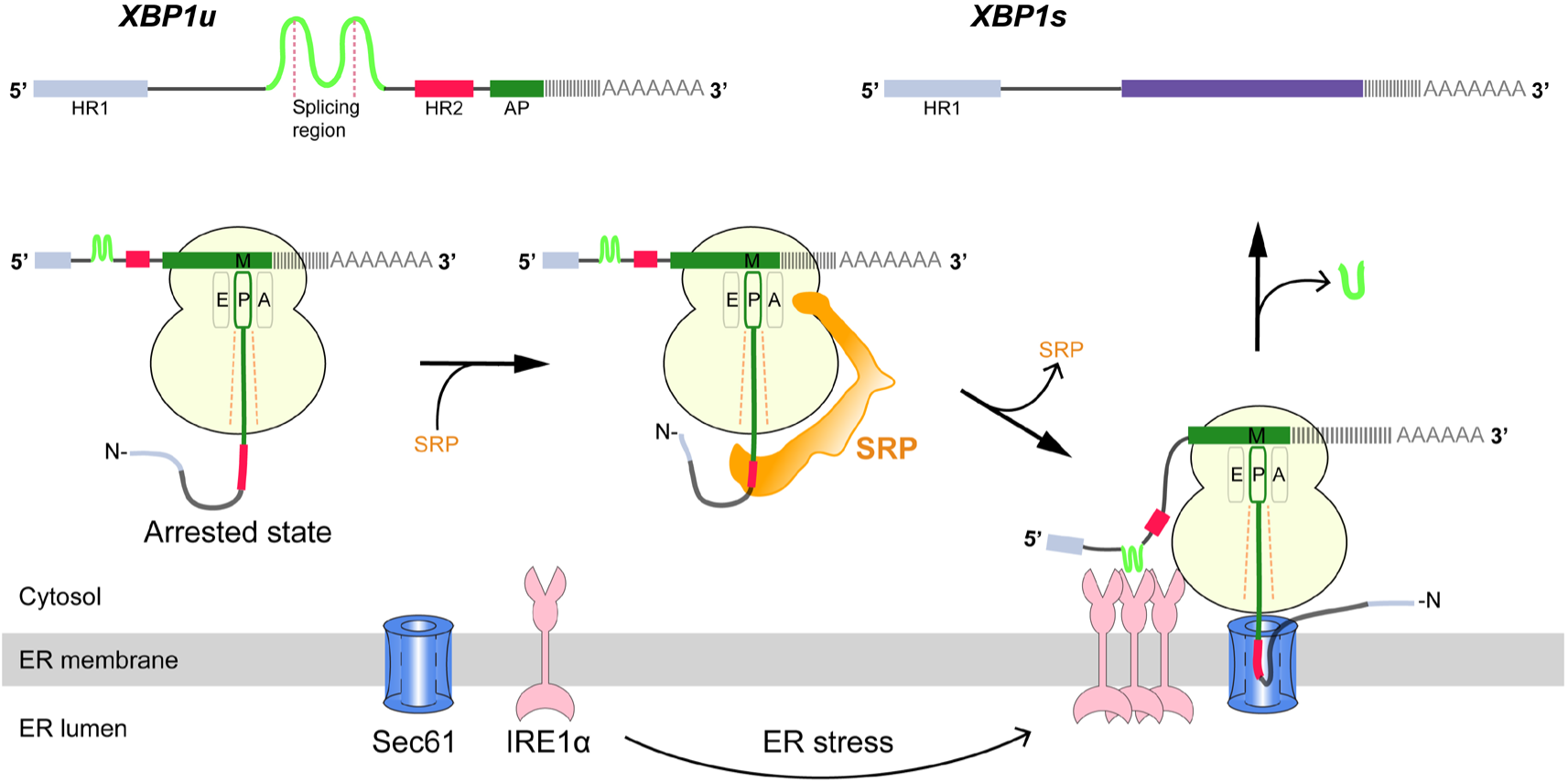
Schematic representation of the IRE1α-XBP1u pathway mediating UPR. Interaction of the XBP1u nascent chain with the ribosomal exit tunnel leads to translational pausing, resulting in SRP recruitment to the RNC, followed by targeting to Sec61 on the ER membrane. IRE1α localized near Sec61 during ER stress can splice *XBP1u* mRNA to *XBP1s* mRNA, which acts as transcription factor in alleviating ER stress.

IRE1α is also involved in a process called regulated IRE1-dependent decay of mRNA (RIDD), where promiscuous cleavage and therefore inactivation of mRNA by IRE1α during ER stress reduces protein influx to the ER (Hollien, Julie and Weissman, 2006; Hollien et al., 2009). It has been shown that active IRE1α can associate with the Sec61 translocon (Plumb et al., 2015), thereby facilitating its access to mRNAs coding for secretory and membrane proteins.

Cytosolic *XBP1u* mRNA is recruited into the proximity of IRE1α on the ER membrane via an ingenious mechanism (Figure 1). XBP1u has two hydrophobic domains, HR1 and HR2, and a C-terminal translational arrest peptide (AP) of about 26 residues which pauses the translating ribosome when residing in the ribosomal exit tunnel (Yanagitani et al., 2011, 2009). During this temporary pause in translation, HR2 is exposed outside the ribosome exit tunnel and can recruit the signal recognition particle (SRP). As a result, the paused XBP1u ribosome–nascent-chain mRNA complex (XBP1u-RNC) is targeted to the Sec61 protein-conducting channel on ER membrane, where mRNA splicing by IRE1α is now possible (Kanda et al., 2016; Plumb et al., 2015). Given the moderate hydrophobicity of HR2, translational pausing is required for efficient recruitment of SRP by the stabilized XBP1u-RNC, and is critical for proper IRE1α-mediated UPR (Kanda et al., 2016; Plumb et al., 2015).

Many studies have focused on the role of IRE1α in the UPR, but the molecular underpinnings of XBP1u-induced translational pausing have not been defined. Here, we have used two complementary approaches, structural analysis and saturation mutagenesis, in order to decipher the structural basis and mechanism of the XBP1u AP activity. We show that the XBP1u AP forms a unique turn in the upper part of the tunnel and makes extensive contacts with ribosomal tunnel components. Notably, the conformation of the XBP1u AP is unaltered within the ribosomal tunnel when the paused complex is bound to SRP or to the Sec61 complex, implying that the XBP1u AP does not function as a force-sensitive switch in the UPR pathway *in vivo*. By saturation mutagenesis, we observe that most but not all XBP1u residues constituting the turn are optimized for translational arrest. Finally, we identify XBP1u AP variants of increased arrest potency, which may be useful as tools for *in vitro* force-sensing studies.

## Results

### Generation and cryo-EM analysis of XBP1u-paused ribosome-nascent chain complexes

We structurally characterized the paused ribosomal complex (XBP1u-RNC) by cryo-EM and single particle analysis using a mutant version of XBP1u (S255A, full length numbering), which was shown previously to pause ribosomes more efficiently than wildtype XBP1u (Yanagitani et al., 2011). The construct used for the RNC preparation encompassed only the HR2 domain and the XBP1u pausing sequence denoted as AP, with N- and C-terminal tags for affinity purification and detection purposes (for clarity we number the residues according to their position in the full-length protein), Figure 2A.

**Figure 2.**
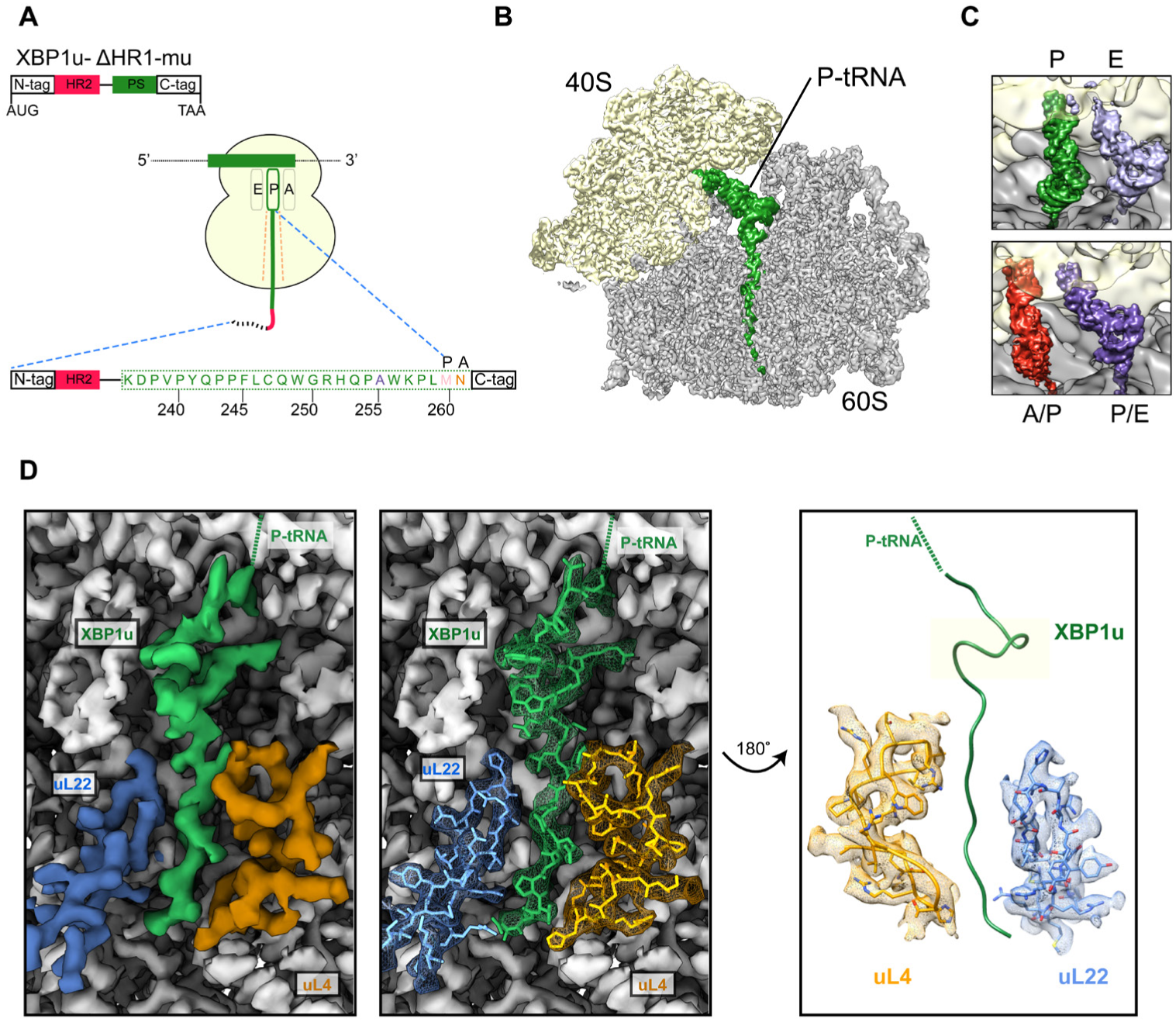
Structural analysis of XBP1u mediated ribosomal pausing. **(A)** Schematic representation of the XBP1u-del-HR1-mu construct used for purification. The construct encodes N-tag, hydrophobic region 2 (red), AP (green) and C-tag. Model for nascent chain in the tunnel, and P-site and A-site positions were denoted as well. **(B)** Traverse section of cryo-EM structure of the paused XBP1u-RNC showing the peptidyl-tRNA (green) with small and large subunits colored in yellow and grey respectively. **(C)** Close-up views showing the two tRNA states of the XBP1u-RNC, post (top panel) and rotated (bottom panel). **(D)** Overview of the XBP1u nascent chain in the ribosomal tunnel. Surface representation of the electron density: P-tRNA (green), uL4 (orange), uL22 (blue) and ribosomal tunnel (grey).

Following translation of the capped *XBP1u* mRNA in a rabbit reticulocyte lysate (RRL) *in vitro* translation system, paused ribosomal complexes were purified using the N-terminal His-tag on XBP1u and subjected to cryo-EM analysis. Processing of the cryo-EM dataset yielded a total of 531,952 ribosomal particles (Figure S1), and multiple rounds of *in silico* sorting for homogenous populations resulted in ~60% of programmed ribosomes (Figure 2B), with the major population of ribosomes in the non-rotated state (~42%, P- and E- site tRNA) and a minor population in the rotated, not yet fully translocated state (~18%, A/P- and P/E- site tRNA, Figure 2C). In both states we observed strong density for the XBP1u chain, which was connected to tRNA and extended down the ribosomal exit tunnel. The average resolutions of the paused complexes were 3.0 Å (Figure S2A) for the post state and 3.1 Å (Figure S2B) for the rotated hybrid state, respectively, with the ribosomal core reaching a resolution of 2.5 Å (Figure S2A). A major portion of the XBP1u peptide in the exit tunnel was resolved to between 3.0 – 3.5 Å for both classes (Figure S3A, B), whereas the resolution in the lower part of the tunnel near the exit was worse than 4 Å, apparently due to flexibility of the nascent chain. We could model 24 amino acid residues of XBP1u, covering the entire AP. In both states, we observed that the ribosomes are paused with Met260 connected to the tRNA in the P-site, in full agreement with findings from ribosome-profiling analysis of mouse embryonic cells (Ingolia et al., 2011). In the following sections, we will refer to the post-state paused RNC complex for further analysis and discussion, since the nascent chain conformation is indistinguishable in both states.

### XBP1u nascent chain in the ribosomal tunnel

The majority of the visible XBP1u nascent chain adopts an extended conformation, except in the upper part of the tunnel where the AP forms a prominent turn (Figure 2D). The turn is comprised of eight residues from W249 to W256, and involves the C-terminal half of the characterized XBP1u AP. Notably, the beginning of the turn is only four residues away from the PTC, suggesting that the turn within the tunnel may be critical for the pausing activity of XBP1u. Of the eight turn-forming residues, six have been previously shown to be critical for pausing by alanine scanning mutagenesis (Yanagitani et al., 2011), c.f., below. Interestingly, residue 255 that has been mutated from Ser to Ala in the sequence used here is part of the turn: A255 is tightly packed in the structure and the larger Ser residue may be sterically more problematic, possibly explaining why the S255A mutation makes the XBP1u a stronger AP.

The turn is located in close proximity to the PTC, above the constriction at uL22 and uL4, the narrowest portion of the tunnel. The conformation of the XBP1u peptide in the lower parts of the tunnel is similar to that observed for a non-pausing mammalian nascent chain in the mammalian ribosome (Voorhees and Hegde, 2015) (Figure S3C – D) and to other known viral and bacterial APs (CMV, MifM and VemP, Figure S3E – G) (Matheisl et al., 2015; Sohmen et al., 2015; Su et al., 2017). However, the turn observed for XBP1u is unique, and is located in a part of the tunnel where some other APs adopt α-helical secondary structure (Figure S3E – G).

### Interactions stabilizing the XBP1u peptide conformation

The turn in the XBP1u AP makes several key interactions with the tunnel wall and is in part in close proximity to the PTC. It is framed by two tryptophan residues (W249 & W256) and protrudes into a hydrophobic crevice in the tunnel, causing the displacement of the base G3904 (Figure 3E). The corresponding base in prokaryotes, A2058, constitutes, together with A2059, the so-called A-stretch in the *E. coli* ribosome which is critical for macrolide binding and drug-mediated ribosome stalling (Wilson, 2009). Moreover, it has been implicated in directly sensing the presence of the nascent peptide in the exit tunnel. Therefore, it is possible that this region evolved also in eukaryotes to contribute to the sensing of nascent chains in the tunnel. The positively charged side chain of Arg251 in XBP1u forms a stabilizing salt bridge with the phosphate of A4388 (Figure 3F), whereas Gly250 and Gln253 engage in hydrogen bonds with A3908 and U4555, respectively (Figure 3G, I). Finally, Trp249 stacks internally onto Gln248 of XBP1u, and the backbone carbonyl of Arg251 makes a hydrogen bond to Lys257 within the XBP1u nascent chain (Figure 3A, 3B). Lys257 also stacks onto U4532, and this stacking interaction might influence the movement of the critical PTC base U4531 (U2585 in *E. coli*) (Figure 3D). Taken together, five of the eight residues that constitute the turn engage in contacts with the tunnel.

**Figure 3.**
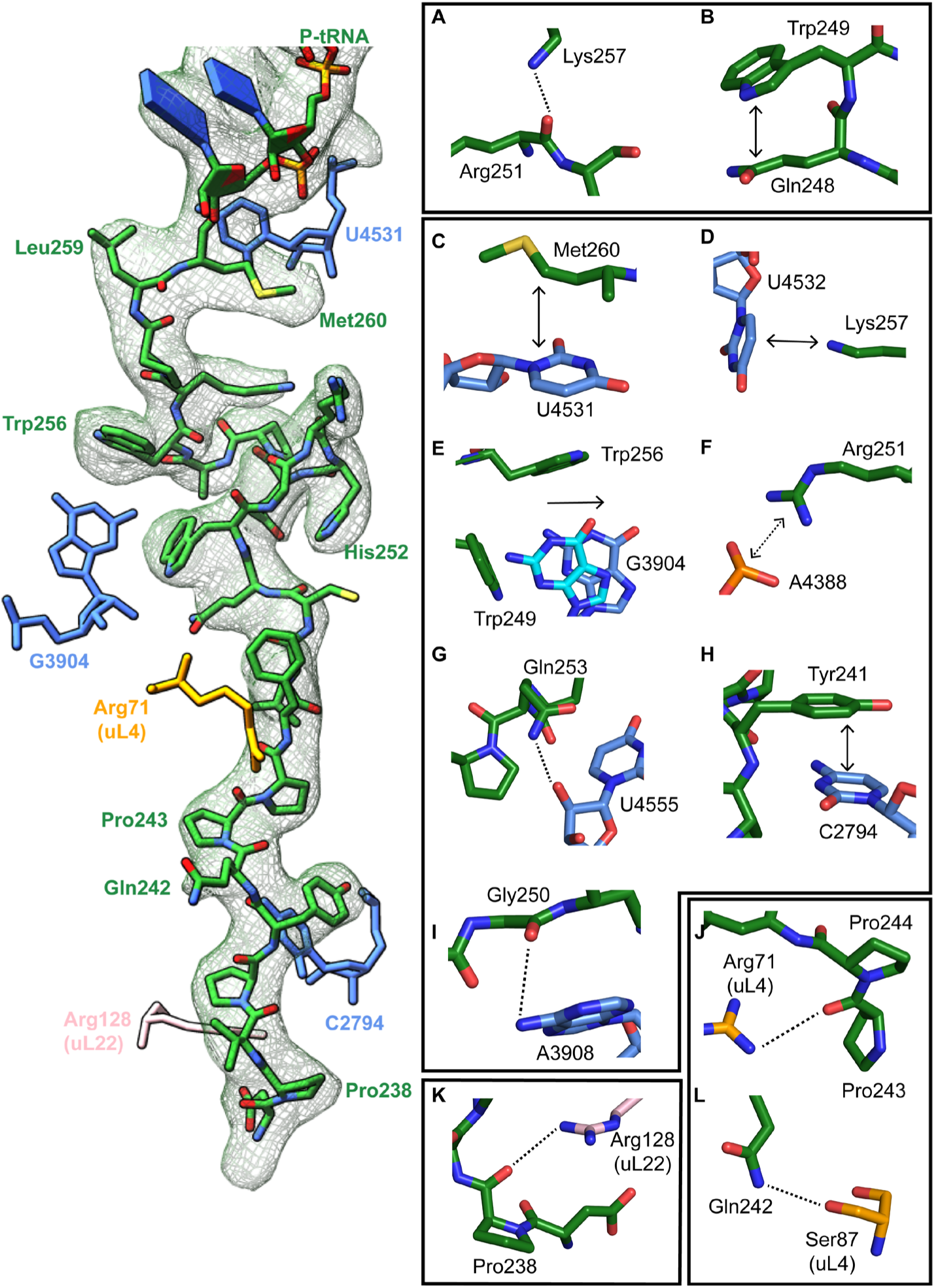
Stabilizing interactions of XBP1u nascent chain with the ribosomal exit tunnel. On the left shown nascent chain model (green) with density (grey mesh), and some interacting 28S rRNA bases and ribosomal protein residues are shown. **(A)** Lys257 of XBP1u (green) is at the hydrogen bond making distance internally within XBP1u residue Arg251. **(B)** Trp249 of the XBP1u stacks internally onto Gln248. **(C)** Met260 of XBP1u makes a hydrophobic interaction with U4531 of 28S rRNA (blue). **(D)** Lys257 stacking with the base U4532 **(E)** Trp256 and Trp249 of XBP1u displace a ribosomal tunnel base G3904 (blue). G3904 conformation with XBP1u is compared with didemnin B treated ribosome (cyan, PDB ID 5LZS). **(F)** Arg251 of XBP1u makes a salt-bridge interaction with the exit tunnel base A4388. **(I, G)** Gly250 and Gln253 of XBP1u are in the distance for making hydrogen bond interaction with 28s rRNA bases A3908 and U4555. **(H)** Tyr241 of XBP1u stacks onto C2794. **(J-L)** Constriction site protein residues making interaction with XBP1u are shown: uL4 (orange) and uL22 (pink).

In the lower part of the tunnel, Tyr241 of XBP1u stacks with C2794 of 28S rRNA (Figure 3H). Three of the remaining interactions of the nascent chain in the lower tunnel region are mediated by the constriction proteins uL4 and uL22, respectively. Here, Arg71 and Ser87 of uL4, as well as Arg128 of uL22 make contacts mostly with the backbone of the nascent chain (Figure 3J – L).

### PTC silencing by the XBP1u peptide

Next, we asked how the unique conformation of the XBP1u peptide in the tunnel results in silencing of the peptidyltransferase activity to cause ribosomal pausing. To that end, we compared the observed PTC conformation with the available mammalian and yeast ribosome structures, either with or without accommodated A-site tRNA, respectively. Since the XBP1u-stalled RNC carries P- and E-site tRNAs but has an empty A-site, we first compared it to the reconstruction of a human 80S ribosome in the post state without A-site tRNA (Behrmann et al., 2015) and of a rabbit 80S ribosome in a pre-accommodation state trapped by didemnin B treatment (Shao et al., 2016) (Figure S4). Both 80S ribosomes display the classical uninduced state of the PTC before full accommodation of tRNA in the A-site, first described for bacterial ribosomes (Martin Schmeing et al., 2005). It is characterized by U4531 (U2585 in *E. coli*) in a typical upward conformation, and C4398 (C2452 in *E. coli*), which is part of the so-called A-site crevice (Hansen et al., 2003), in the typical out-position (Figure S4). In contrast to some bacterial APs, we observed U4531 (U2585) in the XBP1u-stalled RNC in its canonical upward conformation. Although it interacts with the side chain of Met260 (Figure 3C), it appears that this base would not be hindered to switch downwards upon A-site accommodation to adopt the induced conformation. However, C4398 (C2452) is in the closed conformation (Figure 4A, S4), a position that under normal conditions is observed only after A-site accommodation, as in the reconstruction of the yeast 80S ribosome with A-, P-site tRNA and eIF5a (PDB 5GAK) (Schmidt et al., 2015). C4398 (C2452) is stabilized in the closed conformation by Leu259, which, in contrast to Met260, cannot be mutated to alanine without almost entirely loosing stalling activity (see below). Notably, these bases have both been implicated in A-site tRNA accommodation and peptidyl transferase activity. Therefore, the premature positioning of C4398 (C2452) in the closed conformation provides a mechanistic explanation for PTC inactivation by inhibition of A-site tRNA accommodation (Figure 4B). Along the same line, in its observed position Leu259, when interacting with C4398 (C2452), would simply clash with the incoming Asn261 tRNA (Figure 4C). Therefore, inhibition or delay of tRNA accommodation into the A-site appears to be the main mechanism for translational pausing by the XBP1u AP. This idea is further supported by the observation that we do not find a stable class of ribosomes in our population of stalled RNCs that carry a canonical A-site tRNA. Moreover, it can be easily imagined how pulling force applied to the nascent chain can rectify the only mildly perturbed geometry of the PTC and thereby alleviate stalling. Taken together, the entire XBP1u AP contributes to pausing by interacting with the tunnel to form a unique turn structure and, facilitated by this structure, stabilizing the PTC in a conformation that disfavors A-site accommodation.

**Figure 4.**
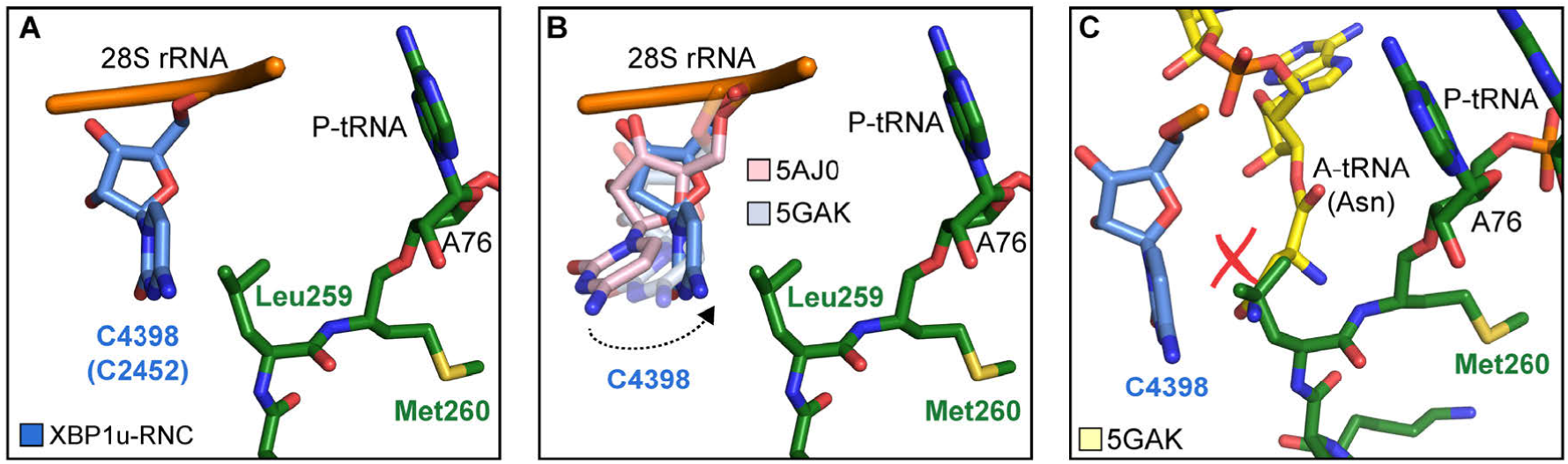
Silencing of peptidyl transferase activity by XBP1u nascent chain. **(A)** Conformation of C4398 in XBP1u-RNC (blue). **(B)** C4398 conformation in the paused complex in comparison with A-site accommodated 80S (PDB ID 5GAK, softblue) and with a post state 80S without an A-site tRNA (PDB ID 5AJ0, softpink) **(C)** Model of an incoming A-site tRNA (yellow, PDB ID 5GAK) clashes with Leu259 of XBP1u. Accommodation of A-site tRNA is prevented by XBP1u.

### Cryo-EM structure of XBP1u-RNC engaged with SRP and Sec61

The paused XBP1u-RNC complex has to be co-translationally targeted to and localized on Sec61 at the ER via the SRP pathway for efficient IRE1α mediated splicing of the XBP1u mRNA (Kanda et al., 2016; Plumb et al., 2015). Due to the AP-triggered prolonged dwell time on the ribosome, the HR2 domain of XBP1u gains sufficient affinity to be recognized by SRP. In order to analyze this special mode of SRP recruitment, and to study the state of the nascent chain within the tunnel when engaged by SRP, we generated cryo-EM structures of the paused XBP1u-RNC complex reconstituted *in vitro* with mammalian SRP or the Sec61 complex.

We reconstituted the purified paused XBP1u-RNC with dog SRP *in vitro* (see Methods for details) and subjected the sample to cryo-EM analysis. After sorting for the presence of SRP and further refinement, a final reconstruction was obtained representing the paused XBP1u-RNC in the post state bound to SRP. We found the characteristic L-shaped density of SRP with its Alu-domain bound to the subunit interface connecting to the S-domain at the exit tunnel (Figure 5A). The final reconstruction had an average resolution of 3.7 Å (Figure S2C) and the SRP itself was resolved between 5 – 10 Å (Figure S5D). A recently published engaged SRP model (PDB 3JAJ) (Voorhees and Hegde, 2015) fits well with our observed density, and individual segments were manually inspected and fitted as rigid bodies in Coot. Analysis of the hydrophobic groove of the SRP54 M-domain, which is known to mediate the recognition and binding of canonical signal sequences, revealed a clear rod-like density resembling that of a signal sequence (Figure 5B). Since the only sufficiently hydrophobic peptide stretch available is HR2 of XBP1u, it is highly likely that this density indeed represents the SRP-bound HR2 domain, bound in a conformation indistinguishable from that of normal SRP-bound signal sequences. Hence, we conclude that the exposed HR2 domain on the paused XBP1u-RNC forms a helical structure upon successful SRP recruitment, which makes a canonical interaction with the M-domain of SRP54. Notably, the nascent chain density was sufficiently well resolved within the tunnel of the XBP1u-RNC-SRP complex to allow for molecular model building. At the given resolution, the conformation of the AP is identical in the presence of SRP to that of the RNC alone. The finding that SRP binding to paused XBP1u-RNCs does not lead to perturbation of the nascent chain within the tunnel strongly suggests that this state maintains the RNC in the paused state.

**Figure 5.**
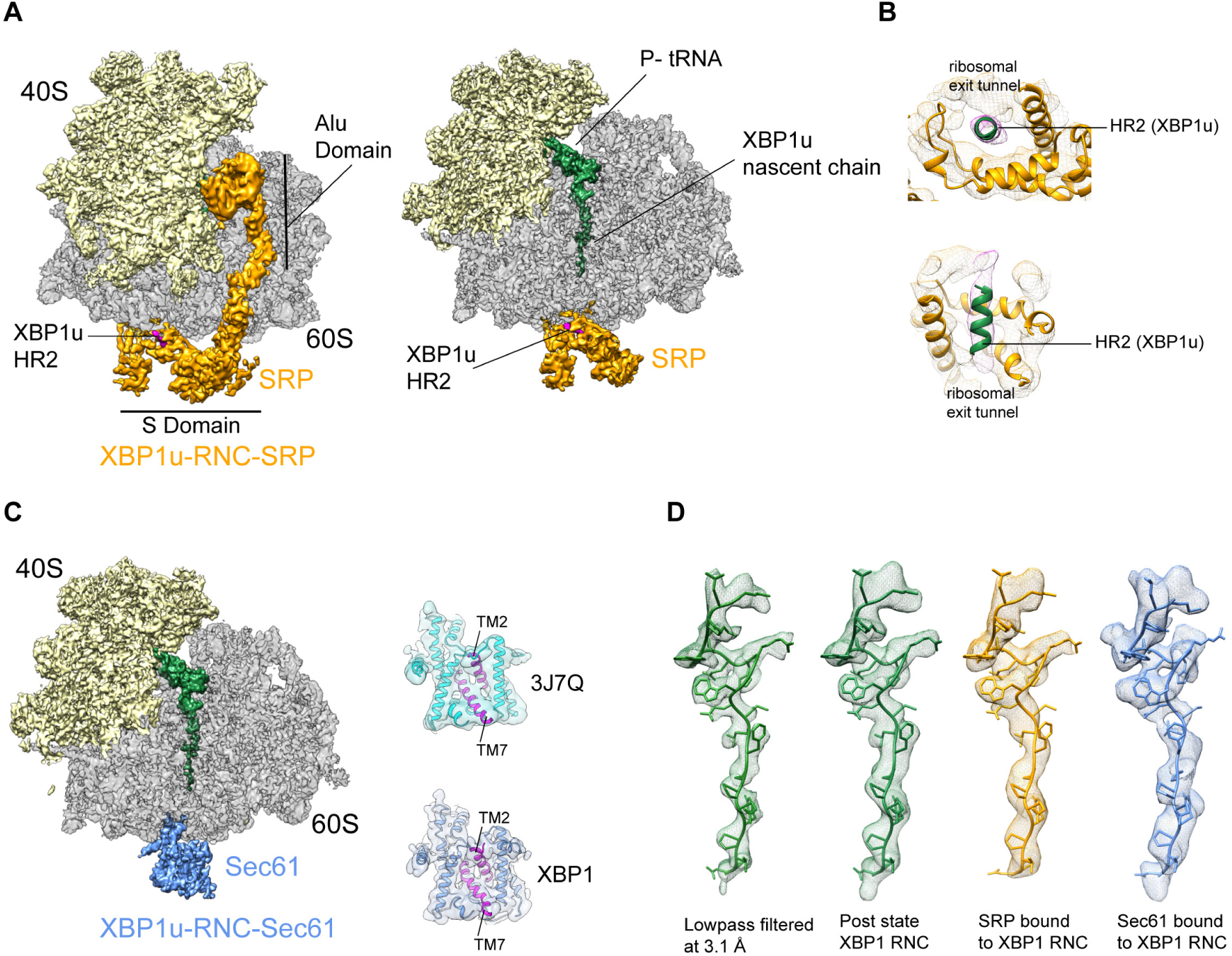
Cryo-EM structures of XBP1u-RNC with SRP and Sec61. **(A)** Cryo-EM reconstruction of XBP1u-RNC with SRP: small (yellow), large (grey), SRP (orange) and hydrophobic region 2 of XBP1u (purple). Same view, a traverse section is shown with XBP1u nascent chain and P-site tRNA (green). **(B)** Close-up view of SRP54 M-domain with a top and cross sectional view showing HR2 of XBP1u. **(C)** Sec61 bound to paused XBP1u-RNC. Cross sectional view: Sec61 (blue), small and large ribosomal subunits, and nascent chain density shown. Idle Sec61 model (cyan, PDB ID 3J7Q) and Sec61 model bound to XBP1u-RNC (blue). Lateral gate is highlighted in both models (purple). **(D)** Unaltered nascent chain in three different states: RNC alone (green), RNC with SRP (orange) and RNC with Sec61 (blue). Density of the nascent chain also colored respectively.

Next, we reconstituted the purified XBP1u-RNC complex with dog PKRM, thereby allowing the XBP1u-RNC-Sec61 complex to form, which should represent the XBP1u-RNC after targeting to the ER. Cryo-EM analysis after solubilization with digitonin resulted in a complex paused in the post state and indeed bound to Sec61. We observed clear density for the Sec61 translocon at the tunnel exit and for the P-site tRNA-attached to nascent chain in the ribosomal tunnel. The average resolution was 3.9 Å (Figure S2D) and the Sec61 complex resolved to a modest resolution of around 8 Å (Figure S5C), due to flexibility as observed before. We performed flexible fitting of the Sec61 structure based on the position of the transmembrane segments in order to analyze the functional state of the translocon and search for additional density possibly representing the HR2 motif. When comparing with known structures of Sec61, we found that our structure represented the idle state with the lateral gate of the translocon, mainly formed by TM2 and TM7, in a closed conformation (Figure 5C). We could not identify any additional density for the HR2 domain of XBP1u on or near the Sec61 complex, indicating a rather weak or transient interaction. Considering the low hydrophobicity of the HR2 domain and previous biochemical evidence that less than 10% of XBP1u becomes integrated into the ER membrane (Plumb et al., 2015), our data are in full agreement with the idea that HR2 can interact with, but cannot productively engage and gate, the Sec61 translocon.

The structure of the XBP1u nascent chain in the XBP1u-RNC-Sec61-complex is indistinguishable from the structures observed in the XBP1u-RNC and the XBP1u-RNC-SRP complexes, with RMSDs between the structures in the range of about 1 Å (Figure 5D). This finding strongly suggests that there is no change in the pausing efficiency during or after successful targeting to the ER membrane, and XBP1u is therefore unlikely to act as a force-sensitive translational switch in the UPR. Probably the long linker length of 52 amino acids between the HR2 domain and the arrest peptide prevents any potential force applied to HR2 upon interaction with SRP or Sec61 to be transmitted to the XBP1u AP.

### Saturation mutagenesis of the XBP1u pausing motif

With the structure in hand, we further characterized the XBP1u AP by saturation mutagenesis. To this end, we placed the XBP1u AP at a variable distance downstream of a hydrophobic segment (H segment) that can generate a pulling force on the nascent chain during *in vitro* cotranslational insertion into rough microsomal membranes (RMs) (Ismail et al., 2012). The construct is composed of an N-terminal part from *E. coli* leader peptidase (LepB) with two transmembrane segments (TM1, TM2), followed by a 155-residue loop, the H segment, a variable-length linker, the 25-residue long human XBP1u AP (with the S255A mutation), and a 23-residue long C-terminal tail (Figure 6A, S6). An acceptor site for N-linked glycosylation located between TM2 and the H segment gets glycosylated by the luminally disposed oligosaccharyltransferase (OST) in molecules that are properly targeted and inserted into the RMs, Figure 6A, while non-glycosylated molecules are indicative of not properly targeted protein and therefore not subjected to pulling forces generated during membrane insertion of the H segment. Hence, only the glycosylated forms of the arrested and full-length species are used for quantitation.

**Figure 6.**
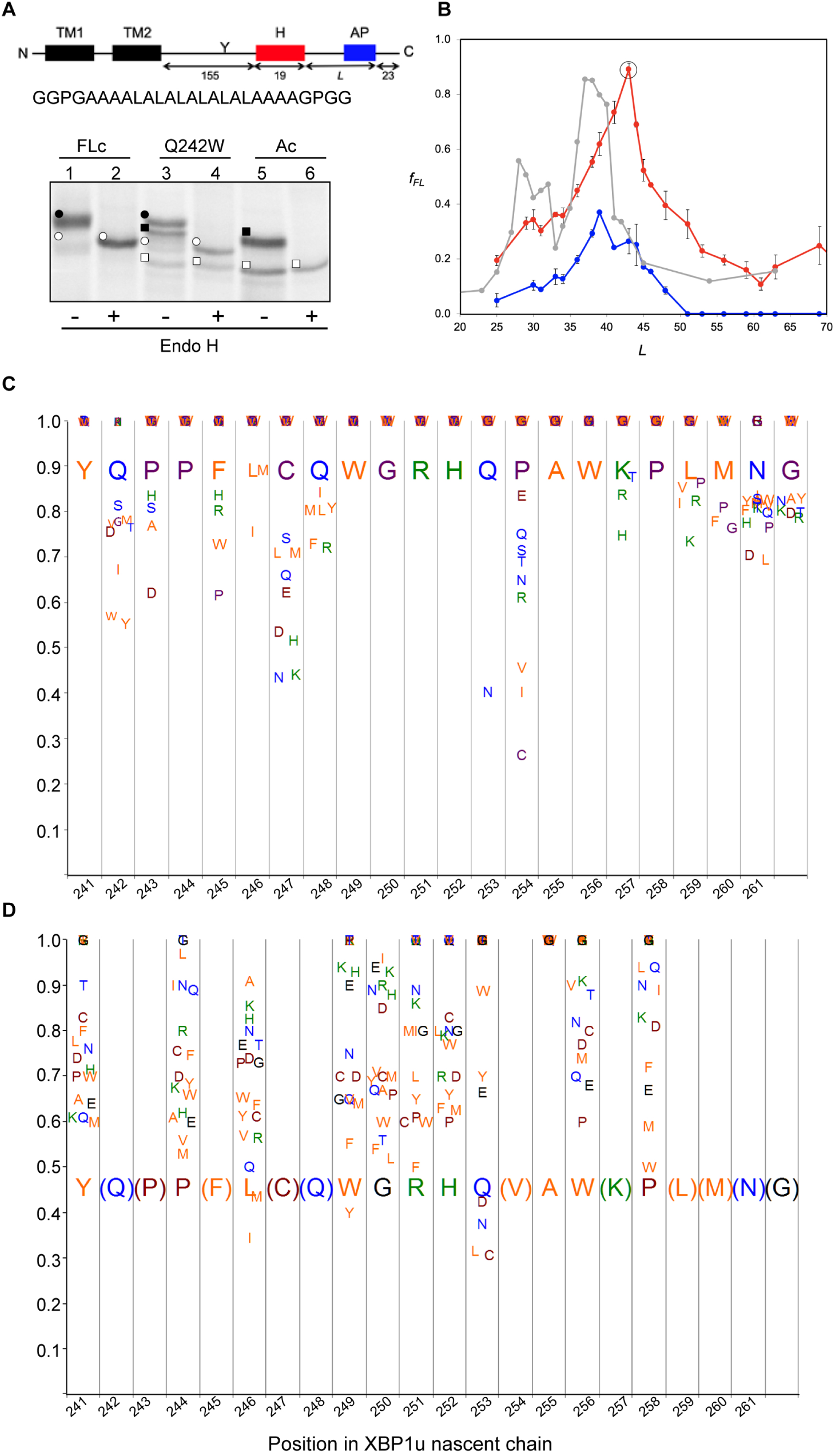
Force profile measurement and saturation mutagenesis of the XBP1u AP. **(A)** Construct used for mutagenesis screens. Y indicates the acceptor site for N-linked glycosylation. The amino acid sequence of the H segment and its flanking GGPG….GPGG residues is shown below. SDS-PAGE gel analysis of a full-length control (FLc, arrest-inactivating mutant), a construct with a Q242W mutation, and an arrest control (Ac) with stop codon immediately downstream of the AP. Full-length species are indicated by circles and arrested species by squares. Black and white colors indicate glycosylated and non-glycosylated species, as shown by Endo H digestion. **(B)** Force profiles measured for LepB-XBP1u (S255A) (red curve) and LepB-XBP1u (S255A, P254C) (blue curve) by *in vitro* translation in RRL supplemented with dog pancreas rough microsomes. A force profile measured in the *E. coli*-derived PURE *in vitro* translation system for the same construct but with the SecM(*Ms*) AP (Ismail et al., 2012) is included for comparison (grey curve). **(C)** Saturation mutagenesis of LepB-XBP1u (S255A, L=43). Residues 241–262 were mutated to all 19 other natural amino acids and *f*_FL_ values were determined. Residues are color-coded as follows: hydrophobic (orange), polar (blue), basic (green), acidic (brown), and G, P and C (purple). **(D)** Same as in *c*, but for LepB-XBP1u (P254V, S255A, L=43).

When a series of constructs with varying linker-lengths is expressed in a rabbit reticulocyte lysate (RRL) *in vitro* translation system supplemented with RMs (Ismail et al., 2012), membrane insertion of the H segment is detected as a peak in a plot of the fraction of full-length protein (*f_FL_*) against the length of the linker+AP segment (*L*, counting from residue N261), Figure 6B (red curve).

Based on this force profile, we chose the construct with maximal pulling force for our initial mutagenesis screen (*L* = 43 residues, compared to *L* = 52 residues between HR2 and the pausing site in the wildtype XBP1u).

Using the LepB-XBP1u[S255A; *L*=43] construct, we systematically changed each of the residues in positions 241 to 262 (position 261 corresponds to the A-site tRNA in the stalled peptide (Ingolia et al., 2011) in the XBP1u AP region to all other amino acids, and measured *f_FL_* for each mutant. The results are summarized in Figure 6C. The majority of the mutations led to weaker arrest (*f_FL_* ≈ 1.0), but a surprisingly large number of mutations reduced *f_FL_* from the starting value of 0.89, indicating stronger arrest variants. Particularly strong reductions in *f_FL_* were seen for mutations P254→[V,I,C], Q253→N, and C247→[N, K], that all have *f_FL_* values < 0.5.

### Structural and mutagenesis hotspots in the XBP1u pausing motif

Some general patterns are discernible from the data in Figure 6C. Many residues in the XBP1u AP are optimal for efficient translational pausing: Y241, P244, W249 to H252, A255, W256, and P258. The turn region in the AP (W249-W256) stands out: six of the eight turn residues are optimal for pausing potency. In contrast, some residues in the AP are clearly sub-optimal in terms of pausing potency: Q242, P243, F245, C247, Q253, P254, K257, and L259.

Mutations in three key residues (C247, Q253 and P254) within the AP lead to particularly strong increases in the pausing strength, with *f_FL_* values in the range 0.2-0.4 (Figure 6C). C247 is located in the lower part of the tunnel, and the introduction of charged or polar residues in this position increases the pausing strength. These residues presumably interact with the ribosomal tunnel by forming hydrogen bonds or salt-bridges with the phosphate backbone of rRNA, but the precise interactions cannot be easily predicted from the structure. Q253 is part of the turn, and when mutated to Asn, the pausing strength is strongly increased. Q253 is positioned in the immediate vicinity of the extremely mutation-sensitive residue A255, and shortening the side chain by one carbon might make the turn better accommodated and more stable in the tunnel. Nine mutations in the neighboring residue P254 also increase the pausing strength, albeit at varying levels. The XBP1u turn is similar to a β-turn, but does not satisfy all the geometrical parameters and therefore is probably less stable than a canonical β-turn. Proline is not favored in β-turns, and its presence in the turn of the XBP1u nascent chain be a result of evolution favoring weaker translational pausing instead of a highly efficient arrest.

We repeated the screen using a stronger version of the pausing motif with the mutation of P254→V from our initial screen (*f_FL_* = 0.46). In this second screen, we focused on positions for which mutations in the first screen gave *f_FL_* ≈ 1, in order to detect any patterns among the mutations that weakened the efficiency of the motif. As can be seen in Figure 6D, all positions except A255 showed a graded response to different mutations; for the latter, all mutations gave *f_FL_* =1.0 (including the back-mutation to the wildtype Ser residue). Interestingly, mutations Q253→ [L, C] led to a reduction in *f_FL_*, despite the fact that the same mutations led to an increase in *f_FL_* in the first screen (Figure 6C). This is most likely due to presence of Val instead of Pro in the neighboring position 254, leading an altered interaction of Q253 with the tunnel or with the nascent chain itself.

We conclude that, although the turn region in the XBP1u AP is nearly optimal for translational pausing, the AP has not evolved to maximize translational arrest potency and considerable stronger versions can be obtained.

### Arrest-enhanced variants of the XBP1u AP can be used as force sensors

Bacterial APs have been used as force sensors to measure forces on a nascent polypeptide chain generated by cotranslational processes such as protein folding or membrane protein insertion into inner membrane (Ismail et al., 2012; Nilsson et al., 2017). To evaluate the possible use of mutant XBP1u APs in such contexts, we re-measured the force profile in Figure 6B using a strong XBP1u AP carrying the mutations S255→A and P254→C (blue curve in Figure 6B). *f_FL_* values are reduced throughout, while the shape of the profile persists. Interestingly, the early peak at *L* ≈ 30 residues seen for the same H-segment constructs expressed in *E. coli* with the SecM AP (Ismail et al., 2012) (grey curve, Figure 6B) is not clearly seen in the mammalian force profiles, suggesting that the H segment interacts differently with the Sec61 and SecYEG translocons at early stages of membrane insertion. Because the mutant AP has a Cys residue in position 254, we considered that the enhanced arrest potency may be due to the formation of a disulfide bond with a ribosomal protein, or within the nascent chain itself. However, no crosslinked product is apparent when a gel is run under non-reducing conditions, Figure S7A, and *f_FL_* is even slightly reduced (as expected from Figure 6C) when the other Cys residue in the AP (C247) is mutated to Ser, Figure S7B.

## Discussion

While a growing number of bacterial APs have been identified, the only reasonably well-characterized mammalian arrest peptide is XBP1u, part of the central regulator in the UPR. We have determined the first high-resolution structure of a mammalian AP stalled in the ribosome exit tunnel and have carried out an extensive mutagenesis analysis, providing insights into its mode of action.

As with previously described APs, XBP1u functions in a unique manner. The XBP1u AP forms a turn within the uppermost part of the tunnel to distort the PTC, inhibiting translational activity. The nascent chain is stabilized within the tunnel and positions the C-terminal region such that the closed conformation of C4398 (C2452) is induced, which is usually adopted only after A-site tRNA accommodation. Together with Leu259, the penultimate amino-acid of the paused nascent chain, this prevents proper accommodation of incoming Asn-tRNA, thereby explaining translational pausing (Figure 7).

**Figure 7.**
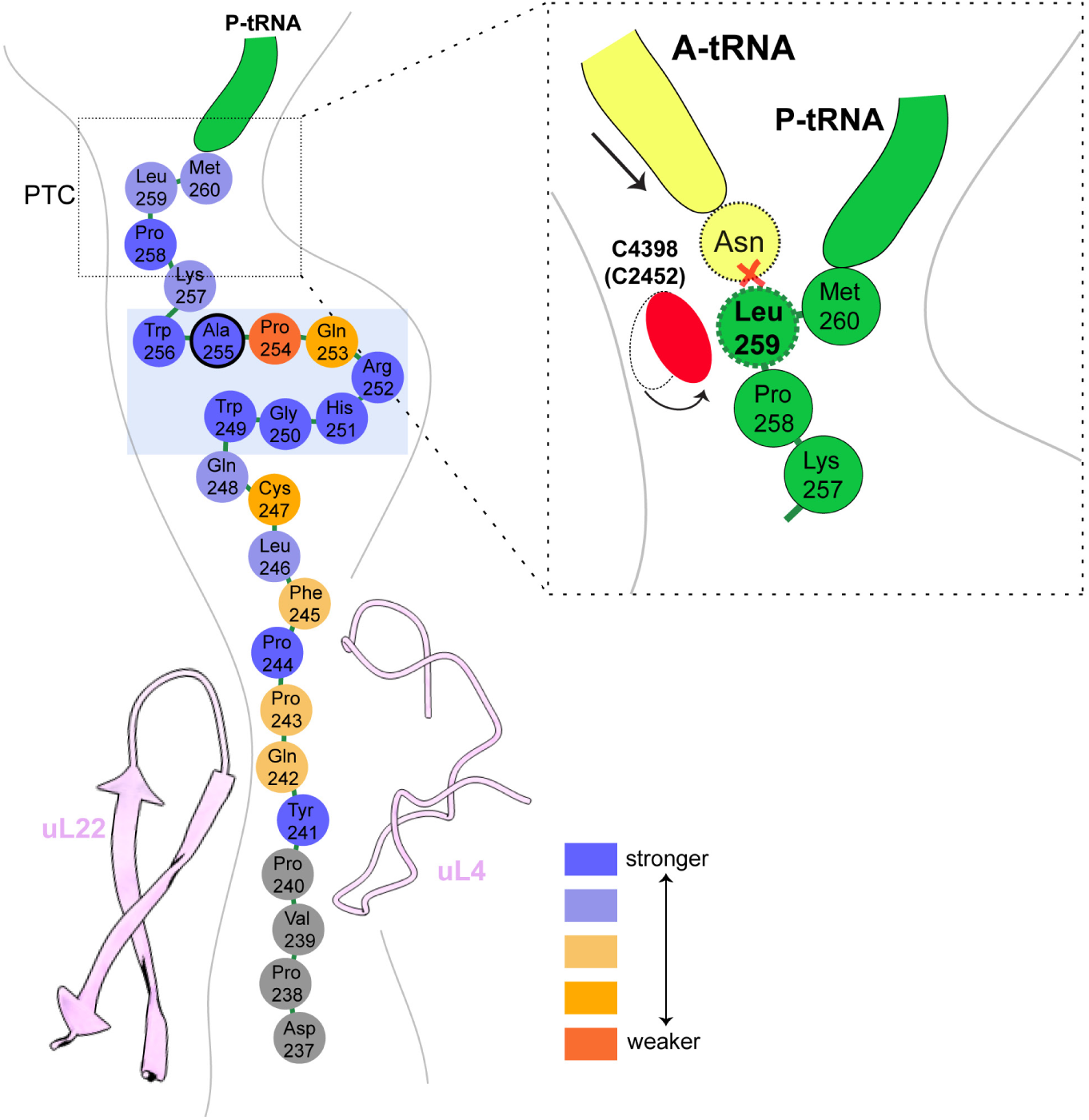
Schematic representation of the XBP1u pausing motif in the exit tunnel. XBP1u residues color coded for pausing potency based on mutagenesis data. Turn formed by XBP1u is highlighted by a light blue box. Inset shows a schematic model of the PTC summarizing the pausing mechanism.

Surprisingly, we show by mutagenesis that the residues constituting the turn in XBP1u have not evolved to maximize ribosome stalling, but rather appear to be selected for an intermediate level of translational arrest potency. P254 plays a critical role in this regard, since nine other residues in this position can all impart stronger arrest potency on the AP. Stronger versions of the XBP1u APs can be useful as force sensors to study cotranslational processes such as membrane-protein insertion into the ER.

We also show that the mildly hydrophobic HR2 segment of XBP1u is recognized as a canonical signal sequence by SRP, with clear density visible in the SRP54 M-domain. However, HR2 cannot engage productively with the Sec61 translocon as a signal sequence, which is consistent with a previous report of minimal membrane insertion observed for XBP1u HR2 (Plumb et al., 2015; Kanda et al., 2016). Finally, we observed that the nascent chain conformation is unaltered within the tunnel in three distinct stages of ER targeting of the ribosome-XBP1u complex: during ribosomal pausing, after recruitment of SRP and upon interaction with Sec61 translocon.

Based on our findings, we propose a structural and mechanistic explanation of XBP1u’s role in the UPR. The XBP1u AP interaction with the ribosomal tunnel pauses ribosomes sufficiently as for the mildly hydrophobic HR2 domain to gain competence for SRP recruitment. The recruitment of SRP ensures proper co-translational targeting, and subsequent localization of the XBP1u mRNA, to the Sec61 translocon on the ER membrane, ensuring efficient cleavage of the *XBP1u* mRNA by IRE1α. The observed unaltered states of the XBP1u nascent chain within the ribosomal tunnel suggest that neither SRP nor Sec61 release the translation stall induced by the XBP1u AP. This is consistent with the previous finding that HR2 is not hydrophobic enough for efficient membrane insertion (Kanda et al., 2016; Plumb et al., 2015). Independently of the nature of this interaction, however, the linker length between the pausing site and the beginning of the HR2 region of XBP1u may also be responsible for uncoupling of HR2 interactions from arrest peptide conformation. From our force profile analysis, the maximal accumulation of full-length product (i.e. maximal force) occurred at a linker length of 43 amino acids, whereas in stalled XBP1u HR2 is around 52 amino acids distant from the PTC, and hence will not exert significant pulling force even if inserted into the ER membrane.

If XBP1u-induced pausing is not released by force, we rather envision two alternative scenarios regarding the fate of the properly targeted, paused complex. First, the pausing may resolve autonomously with the given short half-life or, second, the paused complex is recognized by the Pelota/Hbs1 surveillance system as shown in yeast and recycled. The former is more likely *in vivo*, since the wildtype XBP1u is even weaker than the somewhat stronger mutant (S255A) used in this study. In addition, it has also been shown biochemically that the pause is released when incubated longer during *in vitro* translations (Yanagitani et al., 2011).

In conclusion, the pausing of XBP1u might have evolved as a precise timer, which can pause ribosomes temporarily in order to allow co-translational localization of its polysome-carrying mRNA on the ER membrane for efficient splicing by Ire1α. Interestingly, the mild pausing phenotype is induced by a tight turn of the AP within the exit tunnel, and mirrored by a rather minimal perturbation of the PTC through re-positioning of just one nucleotide, C4398.

## Abbreviations

AP: arrest peptide

## Acknowledgements

This work was supported by grants from the Knut and Alice Wallenberg Foundation (2012.0282), the Swedish Research Council (621-2014-3713), and the Swedish Cancer Foundation (15 0888) to G.v.H., and JSPS KAKENHI JP24228002 to K.K. V.S. was supported by a DFG fellowship through the QBM (Quantitative Biosciences Munich) graduate school. This work was supported by the German Research Council (GRK1721 to R.B.). We also acknowledge the support of a Ph.D. fellowship from the Boehringer Ingelheim Fonds (to K.B.)

## Materials and methods

### Cloning of mutant *XBP1u*

Original full length XBP1u constructs were from the lab of Dr. Kenji Kohno (Nara Institute of Science and Technology, Nara, Japan). The mutant (S255A) full length construct (XBP1u-del-HR1-mu) was then truncated to have only the HR2 domain and pausing motif with N-terminal (8X-His, 3X-Flag and 3C protease cleavage site) and C-terminal (HA-tag) for affinity purification and detection purposes. The final sequence of the construct used for purification:

MGHHHHHHHHGSDYKDHDGDYKDHDIDYKDDDDKDYDIPTTLEVLFQGPGGSISPWILAVLTLQIQSLISCWAFWTTWTQSCSSNALPQSLPAWRSSQRSTQKDPVPYQPPFLCQWGRHQPAWKPLMNYPYDVPDYAGS*

### *In vitro* transcription

The plasmid containing the construct was linearized with Not-I HF enzyme (NEB) at 37°C for 2 h. mRNA for *invitro* translation was prepared using the mMessage mMachine™ T7 Kit (Invitrogen) with linearized plasmid as the template. Capped mRNA was generated following the recommended procedures of the kit. mRNA was then extracted from the reaction mixture using lithium chloride (LiCl) precipitation.

### Rabbit reticulocyte lysate *in vitro* translation

Untreated crude reticulocyte lysate was purchased from Green Hectares (USA), which was then treated with Hemin and MNase, and stored at −80°C. For a 200 µl translation reaction, the 140 µl of treated lysate was used and further supplemented with 3 mM Creatine Phosphate, 30 µM yeast tRNA, 60 mM KOAc, 300 µM Mg(OAc)_2_, 30 µM of amino-acid mixture (Promega) and 0.35 U/µl of RNAse inhibitor (SUPERase. In™, Invitrogen). 80 ng of mRNA per µl of reaction volume was used for subsequent affinity purification of final XBP1u-RNC preparation.

### Purification of XBP1u- ribosome nascent chain complex (XBP1u-RNC)

mRNA was linearized by heating at 65°C for 3 min, before adding it to the translation mixture. 800 µl translation reaction mix was setup and translation was initiated with the addition of capped mRNA. Translation reaction was then incubated in 200 µl aliquots for 10 min at 37°C. Reactions were then stopped by cooling on ice, and then diluted to 2.4 ml with ice-cold buffer A (50 mM HEPES/KOH pH 7.5, 200 mM KOAc, 15 mM Mg(OAc)_2_, 1 mM DTT, 0.1% Nikkol and 0.02 U/µl of RNAse inhibitor). Diluted reaction mix was then incubated with beads at 4°C for 120 min with rotation. After incubation, beads were washed multiple times with buffer A with two intermediate washing steps with buffer A (supplemented with 10 mM imidazole). For elution of the XBP1u- ribosome nascent chain complex, the beads are then incubated with 3C protease (in buffer A) overnight at 4°C with rotation. Flow-through containing XBP1u-RNC were collected, and then centrifuged at 14,000 rpm for 10 min at 4°C to remove any large aggregates. Supernatant from this step was pelleted through 500 mM sucrose (in buffer A) cushion using TLA100.3 rotor at 90,000 rpm for 90 min at 4°C. The preparation yielded 4.2 pmol of XBP1u-RNC which was then used to make cryo-EM grid.

### *In vitro* reconstitution of purified XBP1u-RNC with SRP and Sec61

SRP was purified from a high salt extract of canine rough microsomes by gel filtration (Sephadex G-150), followed by ion-exchange chromatography (DEAE-Sepharose) as described before (B.Martoglio, S.Hauser, 1998). SRP was then further purified by sucrose centrifugation as described before (Walter and Blobel, 1983). XBP1u-RNC-SRP sample is prepared as follows: 1.2X molar excess of purified dog SRP was added to purified XBP1u-RNC in the presence of 2 mM GMP-PNP and 0.01% GDN, and incubated at 25°C for 15 min. Additional 4.5X excess of purified SRP receptor (α and β) and six-fold excess of Sec61 was added and incubated at 25°C for 15 min before being applied onto the grids for cryo-EM analysis.

Canine puromycin/high-salt treated rough membranes (PKRM) were prepared as described before (Gogala et al., 2014). PKRM was pre-treated with RNAsin, and were incubated with purified XBP1u-RNC for 15 min at 25 °C. Membranes were then solubilized with 1.5% digitonin in Buffer A for 90 min in ice. Solubilized ribosome-translocon complexes were pelleted through sucrose cushion (with 500 mM sucrose, 0.3% digitonin, PMSF and protease inhibitor in buffer A). Pelleted complexes were resuspended in buffer A with 0.1% GDN and used for cryo-EM sample preparation.

### Cryo-electron microscopy and single particle reconstruction

XBP1u-RNC (5.2 OD_260_/mL) was applied to 2 nm pre-coated Quantifoil R3/3 grids. Cryo-EM data was collected semi-automatically using EM-TOOLS acquisition software (TVIPS, Germany) on a Titan Krios TEM at a defocus range between 0.5 and 3 µm. All data were recorded on a Falcon II detector (FEI) with a nominal pixel size of 1.084 Å/pixel on the object scale. A total of 6080 micrographs were collected with a total exposure of ~28 electrons/ Å2 fractionated into 10 frames. All micrographs were manually inspected for ice and aggregation, and then subjected to automated particle picking with Gautomatch (https://www.mrc-lmb.cam.ac.uk/kzhang/). All classifications and refinements were performed using Relion-2.1 (Kimanius et al). Total of 531,952 ribosomal particles after 2D classification were subjected 3D classification with a prior round of 3D refinement. Initial 3D classification had two ribosomal states (post and rotated) with tRNA’s. In order to further enrich the post state complex, further 3D classification was done with a mask for P-tRNA and 60S, and the resulting sub-sorted class with 223,773 particles were refined with a masked 60S leading to final resolution of 3 Å. The rotated state from the initial 3D classification with 94,923 particles was also refined with a 60S mask to 3.1 Å overall resolution.

A total of 10,136 micrographs were collected for XBP1u-RNC-SRP dataset and 6,668 were finally subjected to automated particle picking, and further processed as mentioned above. The final sub-sorted class of post state-RNC with SRP was refined with a mask for 60S and SRP.

### Molecular modeling and refinement of the XBP1u-RNC

For the post state XBP1u-RNC, pdb 5LZV (Shao et al., 2016) was used as the initial 80S molecular model of the rabbit 80S ribosome to dock into the sharpened density. The initial fit was done with UCSF Chimera (Pettersen et al., 2004), the model was further adjusted manually in Coot (Emsley and Cowtan, 2004) and refined using phenix.real-space_refine (Adams et al., 2010) with restraints obtained with the command phenix.secondary_structure_restraints. All manual adjustments for the final model were done to fit into corresponding local resolution filtered map generated with Relion 2.1 (Kimanius et al., 2016). Following bases of the 28S rRNA were manually inspected and adjusted in Coot: C2794, G3904, A3908, A4388, C4398, U4531 and U4532. P- and E- tRNA, mRNA was also inspected manually for proper fit into the density.

For the rotated state model, first the large subunit (60S) was fitted. For fitting the 40S, the 40S was split into two parts: the head and the body. Split small subunit models were fitted using Coot and then joined together. P/E- tRNA from the pdb 3J77 (Svidritskiy et al., 2014) and A/P- tRNA from pdb 3JBV (Zhang et al., 2015) were used as initial models in the rotated state.

Refinement for rotated state and XBP1u-RNC with SRP and Sec61 were performed as mentioned above. SRP and Sec61 models were rigid body docked and fitted in Coot, and initial models were from pdb 3JAJ (Voorhees and Hegde, 2015) and 6FTI (Braunger et al., 2018). Molprobity (Chen et al., 2010) was used to calculate the statistics (Table 1) of all the final refined models.

### Enzymes and chemicals

Unless stated otherwise, all chemicals were from Sigma-Aldrich (St Louis, MO, USA). Oligonucleotides were purchased from MWG Biotech AG (Ebersberg, Germany). Pfu Turbo DNA polymerase was purchased from Agilent Technologies. All other enzymes were from Fermentas. The plasmid pGEM-1 and the TNT SP6 Transcription/Translation System were from Promega. [^35^S]Met was from PerkinElmer.

### Construction of mutant library

Site-specific mutagenesis was performed using the QuikChange™ Site-Directed Mutagenesis Kit from Stratagene. All mutants were confirmed by sequencing of plasmid DNA at Eurofins MWG Operon (Ebersberg, Germany).

### Expression *in vitro*

Constructs cloned in pGEM-1 were transcribed and translated in the TNT Quick coupled transcription/translation system. 1 µg of DNA template, 1 µl of [^35^S]-Met (10 µCi; 1 Ci1/437 GBq), 3 µl of zinc acetate dihydrate (5 µM) were mixed with 10 µl of TNT lysate mix, and samples were incubated for 30 min at 30°C. The sample was mixed with 1 µl of RNase I (Affymetrix; 2 mg/ml) and SDS sample buffer and incubated at 30°C for 15 min before loading on a 10% SDS/polyacrylamide gel. Protein bands were visualized in a Fuji FLA-3000 phosphoimager (Fujifilm, Tokyo, Japan). The Image Gauge V 4.23 software (Fujifilm) was used to generate a two-dimensional intensity profile of each gel lane, and the multi-Gaussian fit program from the Qtiplot software package (www.qtiplot.ro) was used to calculate the peak areas of the protein bands. The fraction full-length protein (*f_FL_*) was calculated as *f_FL_* = *I_FL_*/(*I_FL_* + *I_A_*), where *I_FL_* is the intensity of the band corresponding to the full-length protein, and *I_A_* is the intensity of the band corresponding to the arrested form of the protein. Experiments were repeated three times, and SEMs were calculated.

## Supplementary Figures

**Figure S1.**
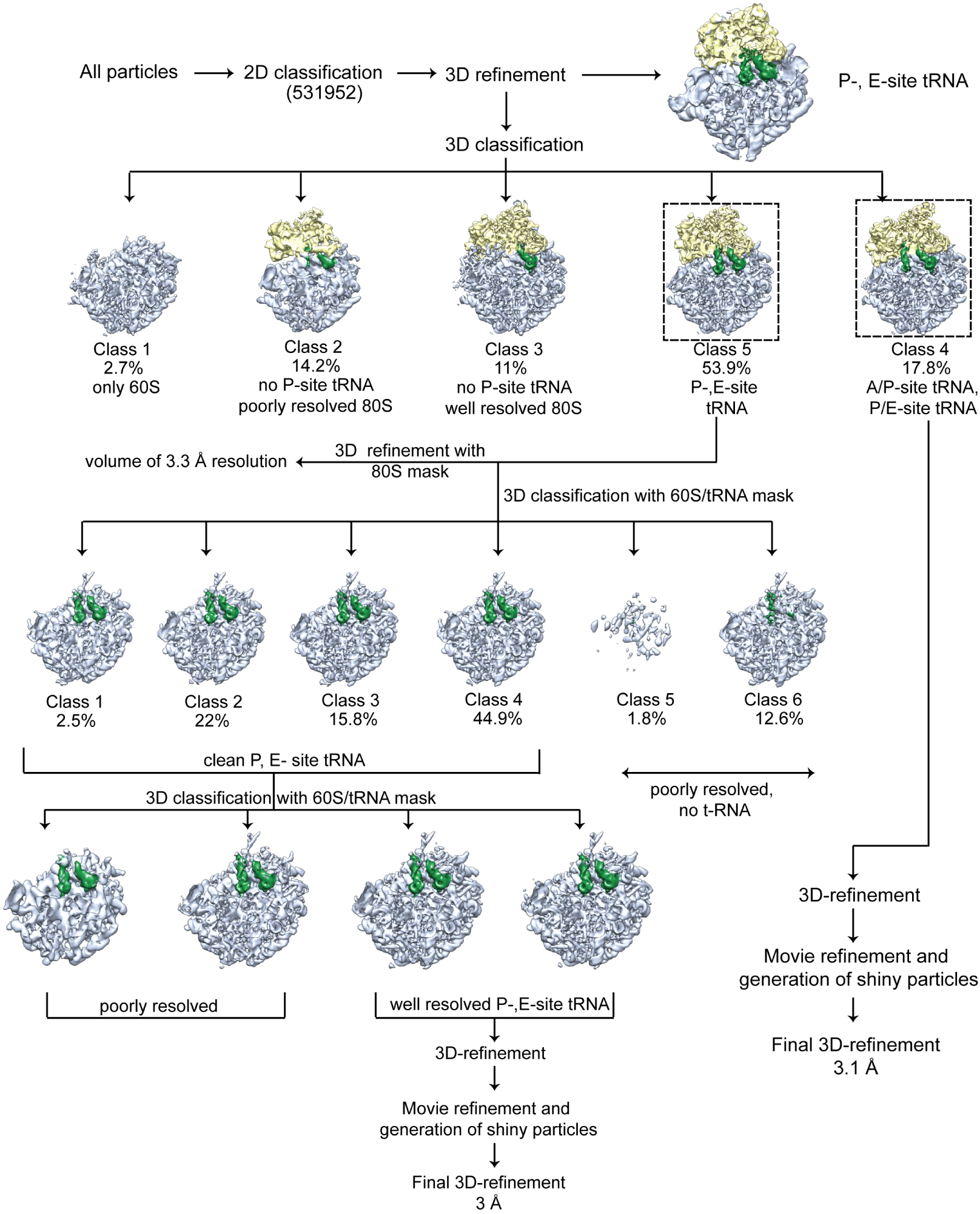
Cryo-EM data processing of the XBP1u nascent chain stalled ribosomes. *In silico* sorting procedure of the data is shown in the schema. Intermediate densities are shown with 60S in blue, 40S in yellow and tRNAs in green. For details check experimental methods section for data processing.

**Figure S2.**
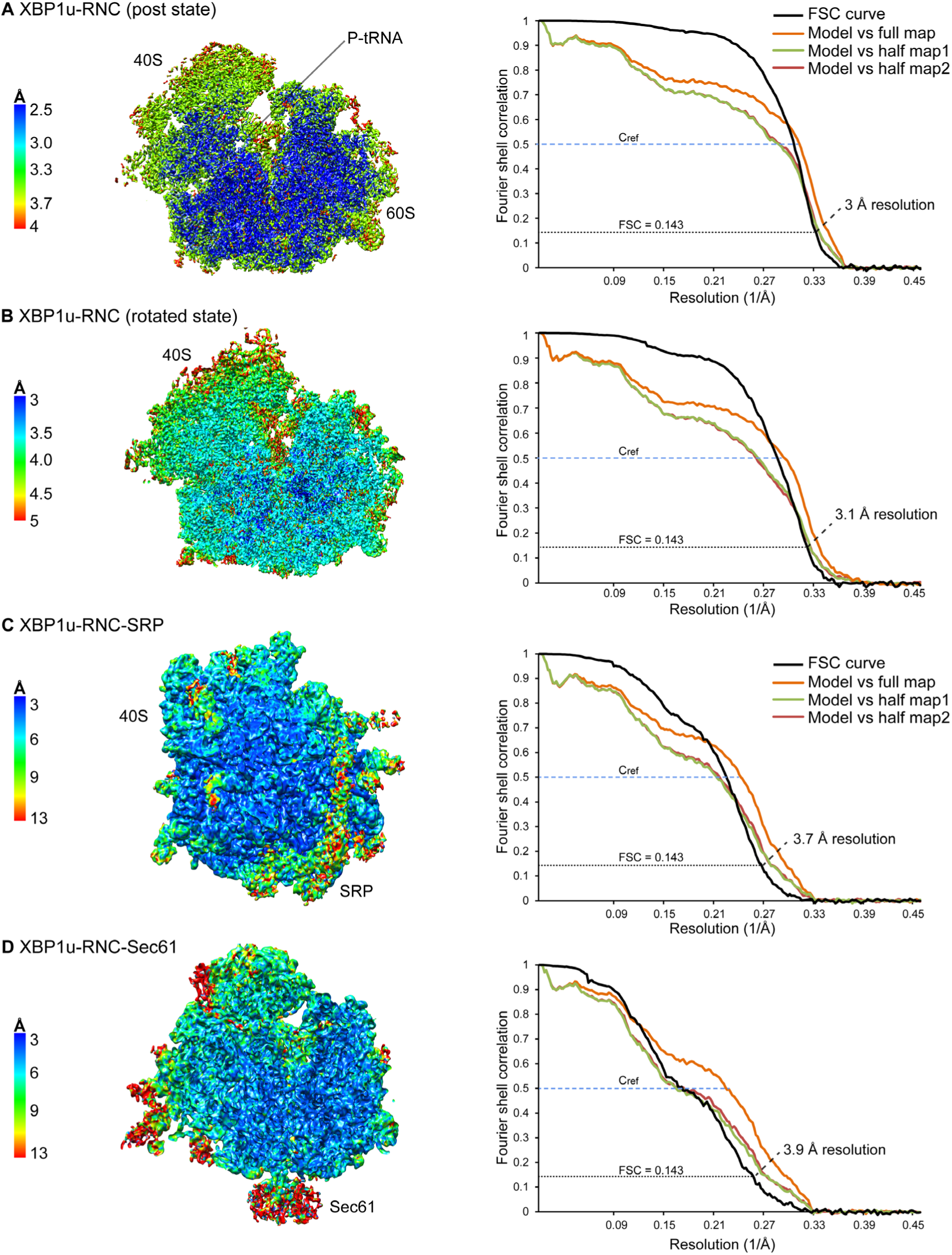
Resolution of XBP1u-RNCs. (Left panel) **(A-B)** Traverse section of post and rotated state of the XBP1u-RNC final map colored according to local resolution are shown here. Relion generated local resolution maps are used. **(C-D)** Electron density maps of XBP1u-RNC with SRP and Sec61 colored according to local resolution. Lowpass filtered maps at 6 Å are used in this figure. Right panel **(A-D)** Fourier shell correlation (black) curve of the final maps, indicating average resolutions (FSC=0.143, dashed black line). FSC curves calculated between final map and model (orange), as well as the self (green) and cross validated (brown) correlation curves for respective XBP1u-RNC states are plotted, indicating resolutions (FSC=0.5 C_ref_, dashed blue line).

**Figure S3.**
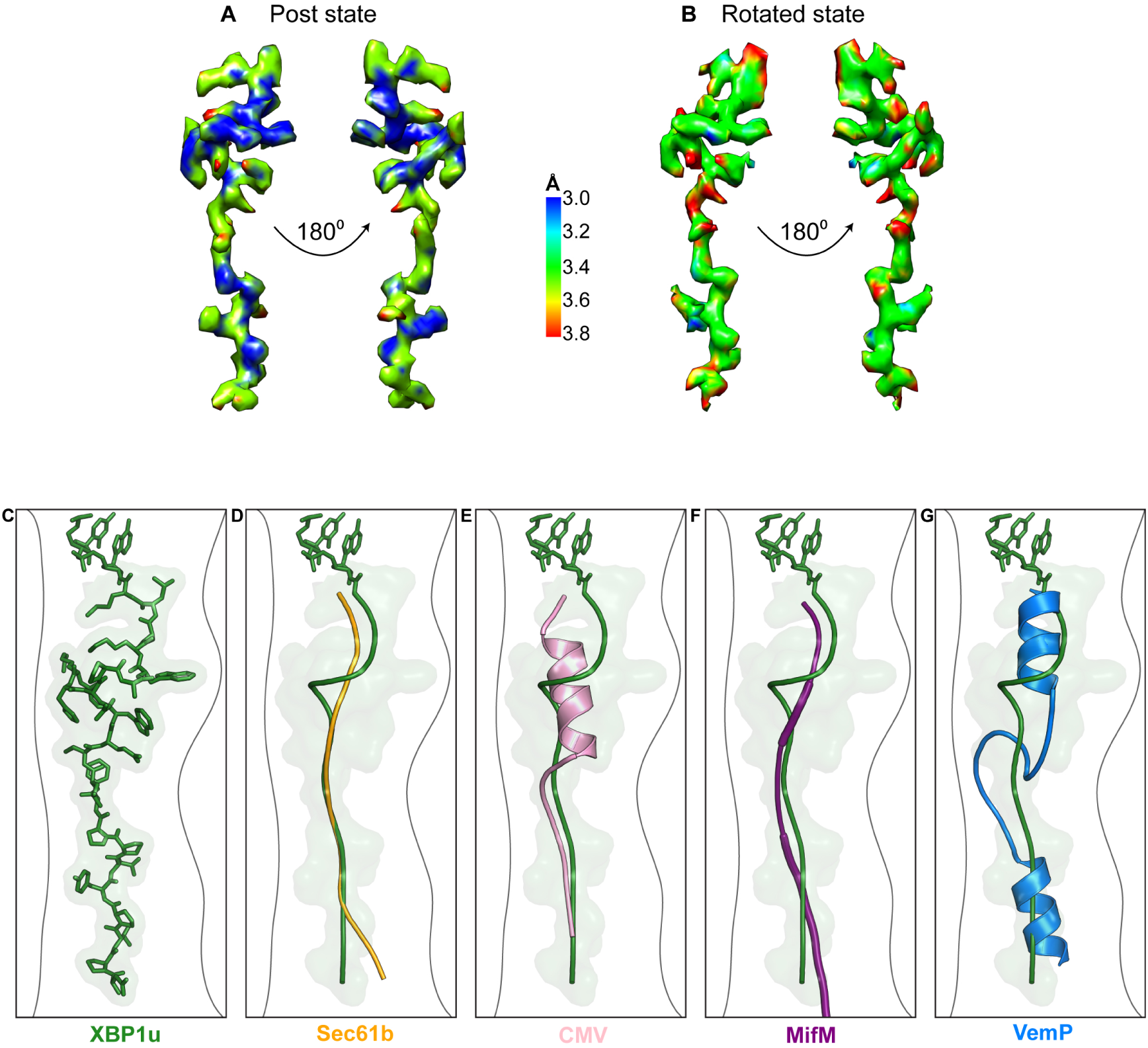
XBP1u nascent chain resolution in the ribosomal tunnel and comparison to other known stalling peptides. **(A-B)** Isolated XBP1u nascent chain density of the post and rotated state XBP1u-RNC colored by local resolution. **(C-G)** Superposition of XBP1u nascent chain (model in forest green, surface in light green) with Sec61b (orange, PDB ID 3JAG) (Voorhees and Hegde, 2015), hCMV (pink, PDB ID 5A8L) (Matheisl et al., 2015), MifM (purple, PDB ID 3J9W) (Sohmen et al., 2015) and VemP (blue, PDB ID 5NWY) (Su et al., 2017) respectively.

**Figure S4.**
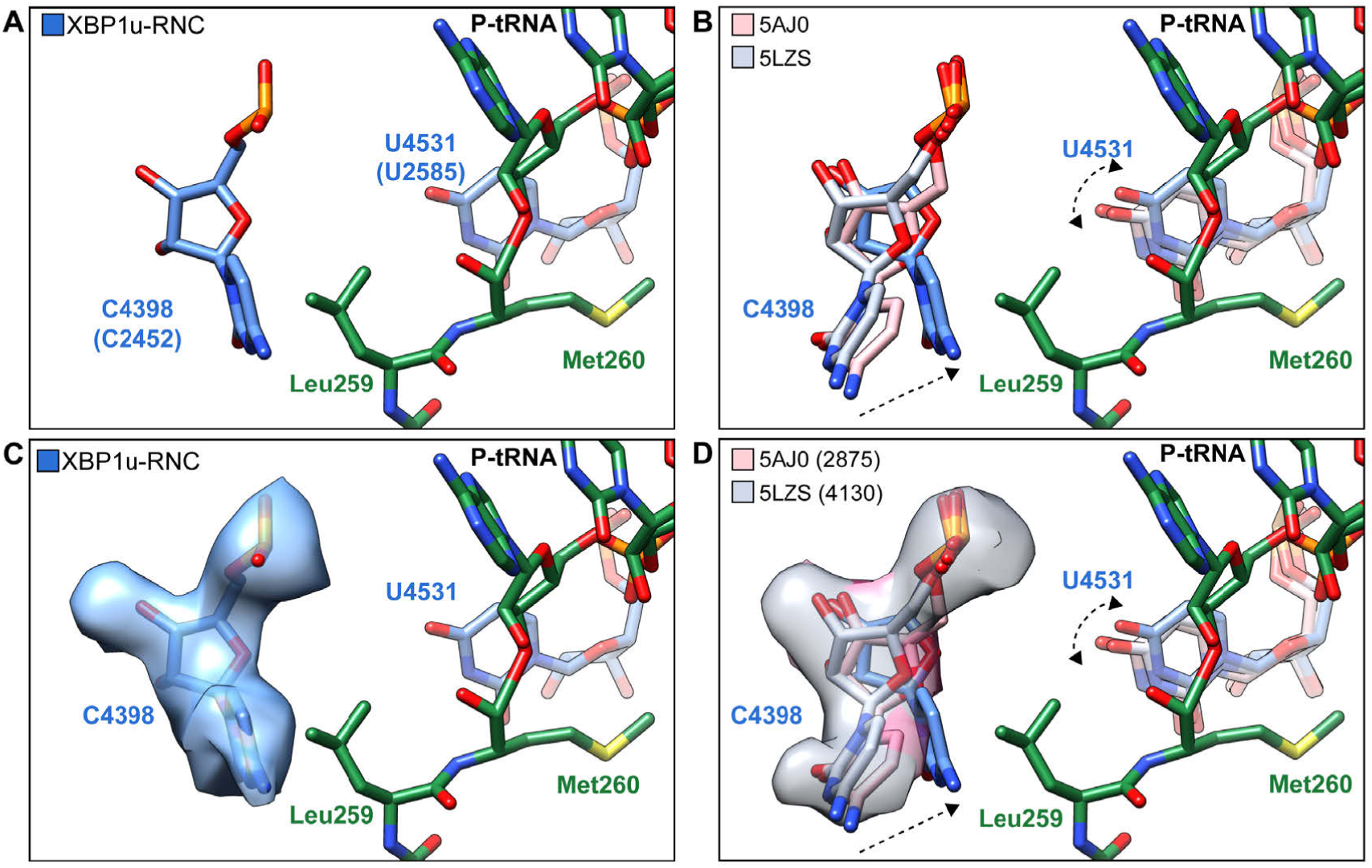
Comparison of C4398(C2452) and U4531(U2585) conformation in XBP1u-RNC with other post-state ribosome 80S models. **(A)** State of the base C4398 and U4531 in XBP1u-RNC (blue). **(B)** State of C4398 and U4531 compared with post state 80S (PDB ID 5AJ0, softpink and 5LZS, softblue) without an accommodated A-site tRNA. **(C)** – **(D)** (A) and (B) displayed with isolated density for the base C4398 (XBP1u-RNC in blue, EMD ID 2875, softpink and 4130, softblue)

**Figure S5.**
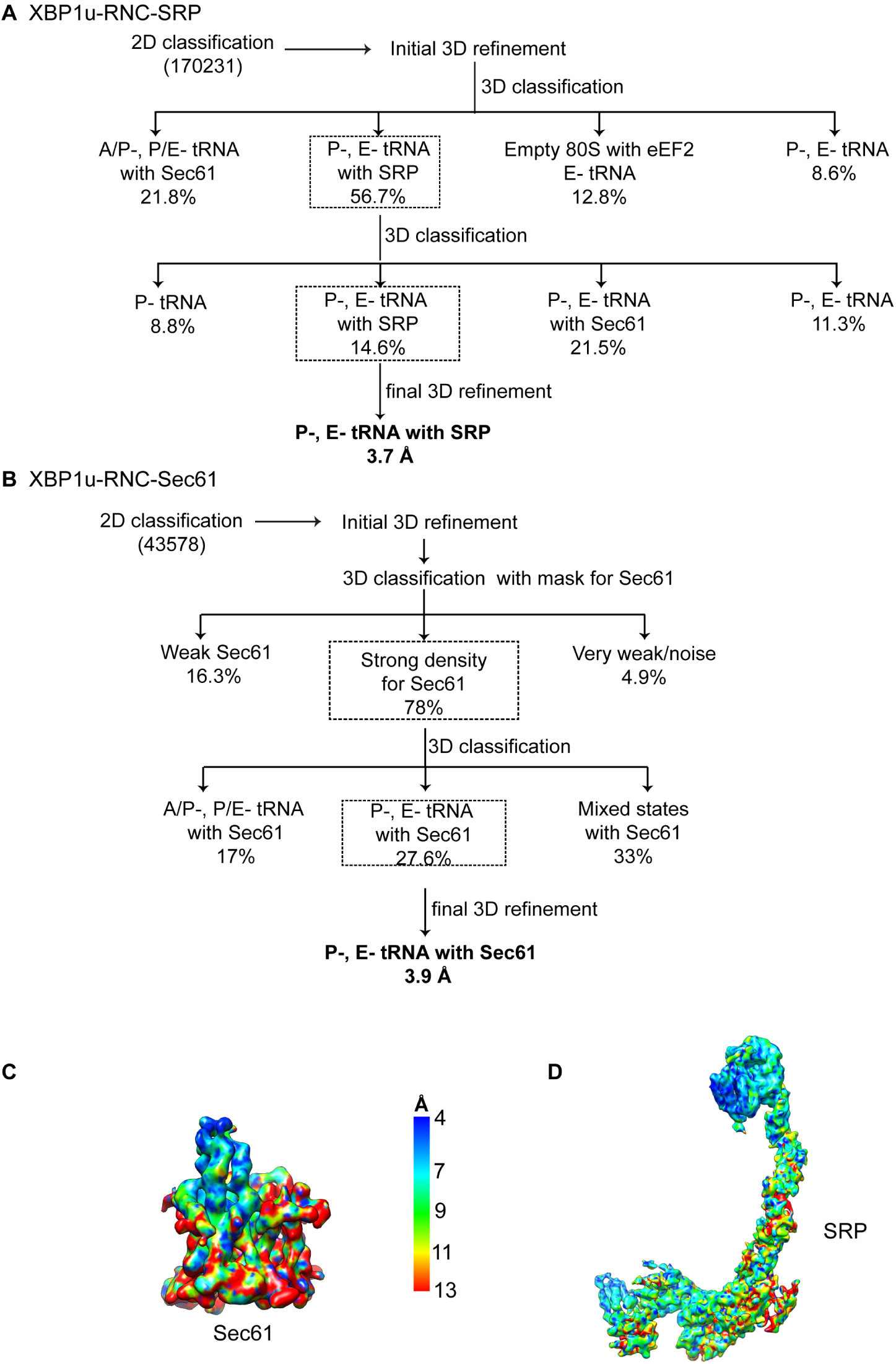
*In silico* sorting and local resolution. **(A-B)** Cryo-EM data processing of XBP1u-RNC-SRP and XBP1u-RNC-Sec61. *In silico* sorting of both the datasets is schematically shown. **(C-D)** Isolated densities of Sec61 and SRP are colored according to their local resolution.

**Figure S6.**
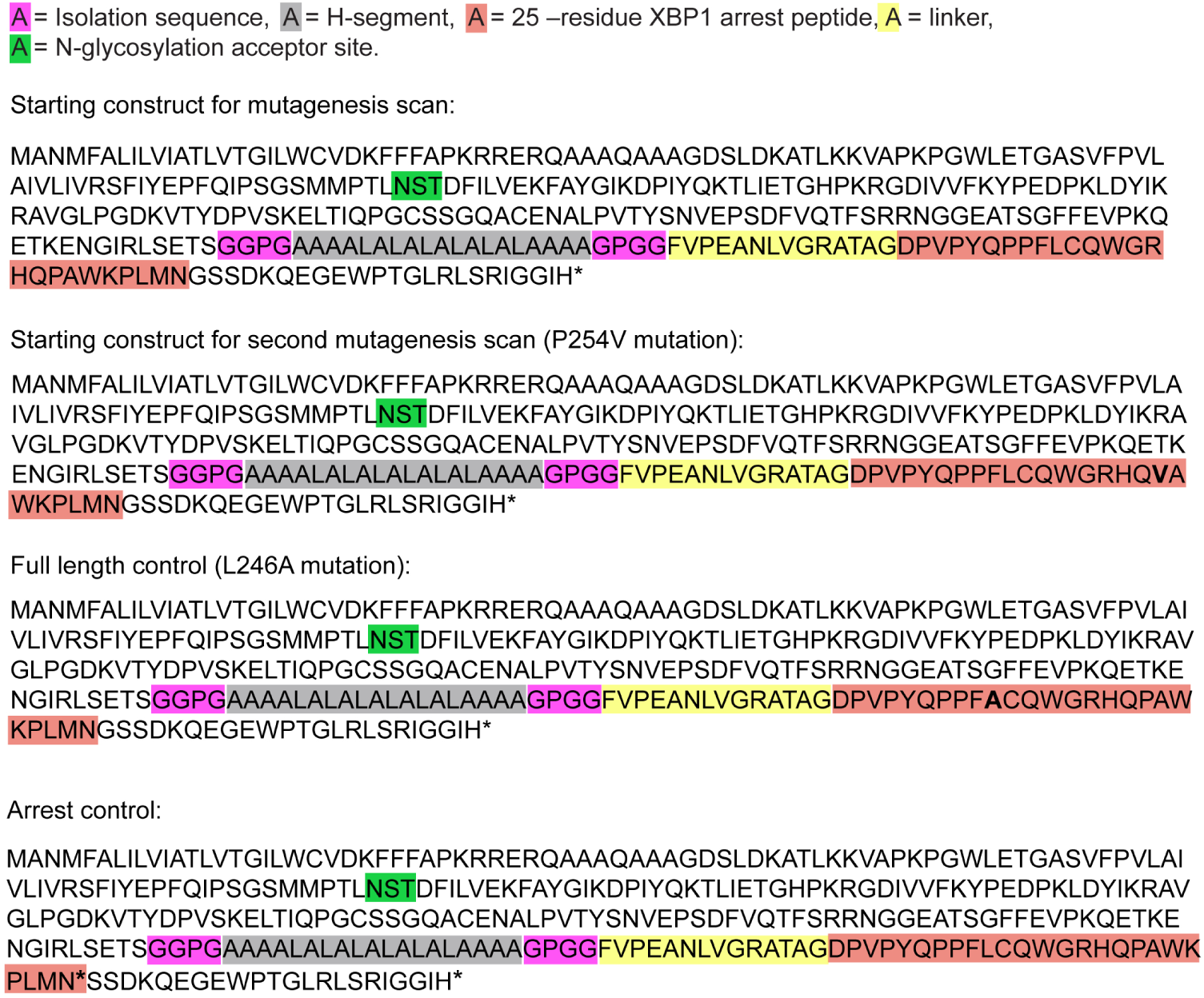
LepB-XBP1u constructs. Amino acid sequences of the LepB-XBP1u[*L*=43] constructs used in the mutagenesis scans.

**Figure S7.**
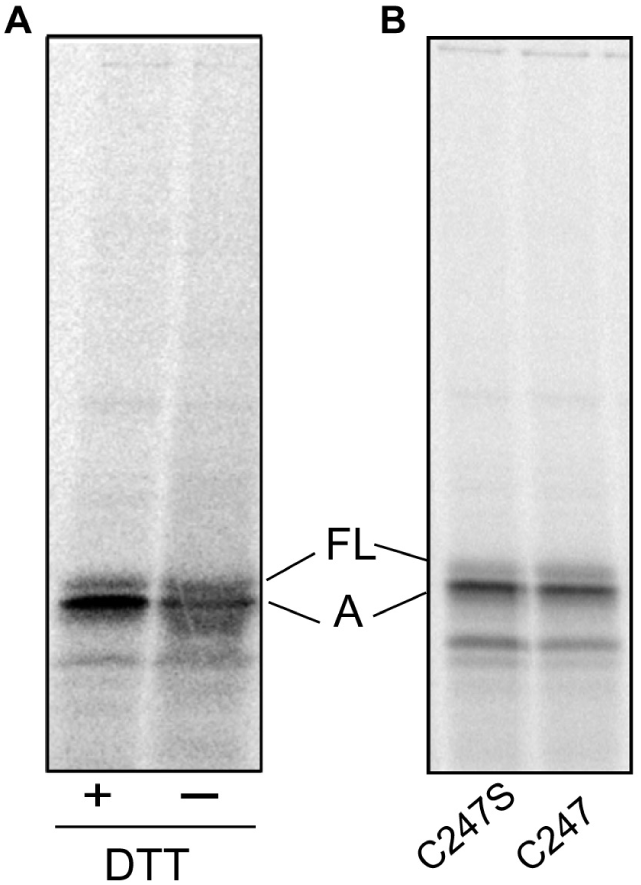
Analysis of Cys positioning by cross-linking. **(A)** No high-Mw disulfide-bonded crosslinked product is seen when *in-vitro* translated LepB-XBP1u[P254C, S255A; *L*=43] construct is analyzed by non-reducing SDS-PAGE in the absence or presence of DTT. **(B)** *f_FL_* is reduced slightly when C247 is mutated to S, from 0.27 for LepB-XBP1u[P254C, S255A; *L*=43] to 0.14 for LepB-XBP1u[C247S, P254C, S255A; *L*=43].

**Supplementary Table 1.** Cryo-EM data collection, refinement and validation statistics. Summary of parameters used during cryo-EM data collection and processing. Refinement and validation statistics are provided for the post state models.

|  | **XBP1-RNC** | **XBP1-RNC** | **XBP1-RNC-SRP** | **XBP1-RNC-Sec61** |
| --- | --- | --- | --- | --- |
| Ribosomal state | Post State | Rotated state | Post state | Post state |
| Microscope | FEI Titan Krios | FEI Titan Krios | FEI Titan Krios | FEI Titan Krios |
| Camera | Falcon II | Falcon II | Falcon II | Falcon II |
| Voltage (kV) | 300 | 300 | 300 | 300 |
| Pixel size (Å) | 1.084 | 1.084 | 1.084 | 1.084 |
| Electron dose (e-/Å<sup>2</sup>) | 28 | 28 | 28 | 28 |
| Defocus range (µm) | 0.5 - 2.5 | 0.5 - 2.5 | 0.5 - 2.5 | 0.5 - 2.5 |
| Particles after 2D (no.) | 531952 | 531952 | 170231 | 43578 |
| Final particles (no.) | 223773 | 94923 | 24875 | 12749 |
| **Model Composition** |  |  |  |  |
| Protein residues | 11717 | 11673 | 12566 | 12239 |
| RNA bases | 5669 | 5797 | 5874 | 5668 |
| **Resolution (Å)** | 3 | 3.1 | 3.7 | 3.9 |
| FSC threshold | 0.143 | 0.143 | 0.143 | 0.143 |
| Map CC (around atoms) | 0.76 | 0.75 | 0.71 | 0.68 |
| Map CC (whole unit cell) | 0.73 | 0.72 | 0.68 | 0.66 |
| Map sharpening B-factor (Å<sup>2</sup>) | -71.2 | -59.9 | -105.54 | -81.6 |
| **RMS Deviations** |  |  |  |  |
| Bond lengths (Å) | 0.004 | 0.0038 | 0.0036 | 0.0035 |
| Bond angles (°) | 0.91 | 0.92 | 0.88 | 0.91 |
| **Validation** |  |  |  |  |
| MolProbity score | 1.5 | 1.66 | 1.55 | 1.5 |
| Clashscore | 4.9 | 4.82 | 5.34 | 4.55 |
| Poor rotamers (%) | 0.23 | 0.20 | 0.16 | 0.13 |
| **Ramachandran Plot** |  |  |  |  |
| Disallowed (%) | 0.03 | 0.09 | 0.05 | 0.02 |
| Allowed (%) | 3.60 | 5.67 | 3.84 | 3.87 |
| Favored (%) | 96.37 | 94.24 | 96.11 | 96.1 |

## References

Adams PD, Afonine P V., Bunkóczi G, Chen VB, Davis IW, Echols N, Headd JJ, Hung LW, Kapral GJ, Grosse-Kunstleve RW, McCoy AJ, Moriarty NW, Oeffner R, Read RJ, Richardson DC, Richardson JS, Terwilliger TC, Zwart PH. 2010. PHENIX: A comprehensive Python-based system for macromolecular structure solution. Acta Crystallogr Sect D Biol Crystallogr 66:213–221. doi:10.1107/S0907444909052925

Aragón T, van Anken E, Pincus D, Serafimova IM, Korennykh A V, Rubio CA, Walter P. 2009. Messenger RNA targeting to endoplasmic reticulum stress signalling sites. Nature 457:736–740.

B.Martoglio, S.Hauser BD. 1998. Cotranslational translocation of proteins into microsomes derived from the rough endoplasmic reticulum of mammalian cells. Cell Biol A Lab Handb 265–273.

Behrmann E, Loerke J, Budkevich T V., Yamamoto K, Schmidt A, Penczek PA, Vos MR, Bürger J, Mielke T, Scheerer P, Spahn CMT. 2015. Structural snapshots of actively translating human ribosomes. Cell 161:845–857. doi:10.1016/j.cell.2015.03.052

Bertolotti A, Zhang Y, Hendershot LM, Harding HP, Ron D. 2000. Dynamic interaction of BiP and ER stress transducers in the unfolded-protein response. Nat Cell Biol 2:326–332.

Braunger K, Pfeffer S, Shrimal S, Gilmore R, Berninghausen O, Mandon EC, Becker T, Förster F, Beckmann R. 2018. Structural basis for coupling protein transport and N-glycosylation at the mammalian endoplasmic reticulum. Science 360:215–219. doi:10.1126/science.aar7899

Calfon M, Zeng H, Urano F, Till JH, Hubbard SR, Harding HP, Clark SG, Ron D. 2002. IRE1 couples endoplasmic reticulum load to secretory capacity by processing the XBP-1 mRNA. Nature 415:92–96. doi:10.1038/415092a

Chen VB, Arendall WB, Headd JJ, Keedy DA, Immormino RM, Kapral GJ, Murray LW, Richardson JS, Richardson DC. 2010. MolProbity: All-atom structure validation for macromolecular crystallography. Acta Crystallogr Sect D Biol Crystallogr 66:12–21. doi:10.1107/S0907444909042073

Credle JJ, Finer-Moore JS, Papa FR, Stroud RM, Walter P. 2005. On the mechanism of sensing unfolded protein in the endoplasmic reticulum. Proc Natl Acad Sci U S A 102:18773–18784. doi:10.1073/pnas.0509487102

Emsley P, Cowtan K. 2004. Coot: Model-building tools for molecular graphics. Acta Crystallogr Sect D Biol Crystallogr 60:2126–2132. doi:10.1107/S0907444904019158

Fagone P, Jackowski S. 2009. Membrane phospholipid synthesis and endoplasmic reticulum function. J Lipid Res 50:S311–S316. doi:10.1194/jlr.R800049-JLR200

Gardner BM, Walter P. 2011. Unfolded Proteins Are Ire1-Activating Ligands That Directly Induce the Unfolded Protein Response. Science 333:1891–1894. doi:10.1126/science.1209126

Gogala M, Becker T, Beatrix B, Armache J-P, Barrio-Garcia C, Berninghausen O, Beckmann R. 2014. Structures of the Sec61 complex engaged in nascent peptide translocation or membrane insertion. Nature 506:107–10. doi:10.1038/nature12950

Görlach A, Klappa P, Kietzmann DT. 2006. The Endoplasmic Reticulum: Folding, Calcium Homeostasis, Signaling, and Redox Control. Antioxid Redox Signal 8:1391–1418. doi:10.1089/ars.2006.8.1391

Grootjans J, Kaser A, Kaufman RJ, Blumberg RS. 2016. The unfolded protein response in immunity and inflammation. Nat Rev Immunol 16:469–484.

Hansen JL, Moore PB, Steitz TA. 2003. Structures of five antibiotics bound at the peptidyl transferase center of the large ribosomal subunit. J Mol Biol 330:1061–1075. doi:10.1016/S0022-2836(03)00668-5

Hollien, Julie and Weissman JS. 2006. Decay of Endoplasmic Reticulum-Localized mRNAs During the Unfolded Protein Response. Science 313: 104–107. doi:10.1080/07373937.2010.483029

Hollien J, Lin JH, Li H, Stevens N, Walter P, Weissman JS. 2009. Regulated Ire1-dependent decay of messenger RNAs in mammalian cells. J Cell Biol 186:323–331. doi:10.1083/jcb.200903014

Ingolia NT, Lareau L, Weissman J. 2011. Ribosome Profiling of Mouse Embryonic Stem Cells Reveales Complexity of Mammalian Proteomes. Cell 147:789–802. doi:10.1016/j.cell.2011.10.002.Ribosome

Ismail N, Hedman R, Schiller N, von Heijne G. 2012. A biphasic pulling force acts on transmembrane helices during translocon-mediated membrane integration. Nat Struct Mol Biol 19:1018–1022. doi:10.1038/nsmb.2376

Kanda S, Yanagitani K, Yokota Y, Esaki Y, Kohno K. 2016. Autonomous translational pausing is required for XBP1u mRNA recruitment to the ER via the SRP pathway. Proc Natl Acad Sci U S A 113:E5886–E5895. doi:10.1073/pnas.1604435113

Kimanius D, Forsberg BO, Scheres SHW, Lindahl E. 2016. Accelerated cryo-EM structure determination with parallelisation using GPUs in RELION-2. Elife 5:1–21. doi:10.7554/eLife.18722

Kimata Y, Ishiwata-Kimata Y, Ito T, Hirata A, Suzuki T, Oikawa D, Takeuchi M, Kohno K. 2007. Two regulatory steps of ER-stress sensor Ire1 involving its cluster formation and interaction with unfolded proteins. J Cell Biol 179:75–86. doi:10.1083/jcb.200704166

Kohno K. 2010. Stress-sensing mechanisms in the unfolded protein response: similarities and differences between yeast and mammals. J Biochem 147:27–33.

Korennykh A V., Egea PF, Korostelev AA, Finer-Moore J, Zhang C, Shokat KM, Stroud RM, Walter P. 2009. The unfolded protein response signals through high-order assembly of Ire1. Nature 457:687–693. doi:10.1038/nature07661

Li H, Korennykh A V, Behrman SL, Walter P. 2010. Mammalian endoplasmic reticulum stress sensor IRE1 signals by dynamic clustering. Proc Natl Acad Sci 107:16113–16118. doi:10.1073/pnas.1010580107

Martin Schmeing T, Huang KS, Strobel SA, Steitz TA. 2005. An induced-fit mechanism to promote peptide bond formation and exclude hydrolysis of peptidyl-tRNA. Nature 438:520–524. doi:10.1038/nature04152

Matheisl S, Berninghausen O, Becker T, Beckmann R. 2015. Structure of a human translation termination complex. Nucleic Acids Res 43:8615–8626. doi:10.1093/nar/gkv909

Mori K. 2009. Signalling pathways in the unfolded protein response: Development from yeast to mammals. J Biochem 146:743–750. doi:10.1093/jb/mvp166

Nilsson OB, Nickson AA, Hollins JJ, Wickles S, Steward A, Beckmann R, Von Heijne G, Clarke J. 2017. Cotranslational folding of spectrin domains via partially structured states. Nat Struct Mol Biol 24:221–225. doi:10.1038/nsmb.3355

Okamura K, Kimata Y, Higashio H, Tsuru A, Kohno K. 2000. Dissociation of Kar2p/BiP from an ER Sensory Molecule, Ire1p, Triggers the Unfolded Protein Response in Yeast. Biochem Biophys Res Commun 279:445–450. doi:https://doi.org/10.1006/bbrc.2000.3987

Pettersen EF, Goddard TD, Huang CC, Couch GS, Greenblatt DM, Meng EC, Ferrin TE. 2004. UCSF Chimera – A visualization system for exploratory research and analysis. J Comput Chem 25:1605–1612. doi:10.1002/jcc.20084

Plumb R, Zhang ZR, Appathurai S, Mariappan M. 2015. A functional link between the co-translational protein translocation pathway and the UPR. Elife 4:2–27. doi:10.7554/eLife.07426

Schmidt C, Becker T, Heuer A, Braunger K, Shanmuganathan V, Pech M, Berninghausen O, Wilson DN, Beckmann R. 2015. Structure of the hypusinylated eukaryotic translation factor eIF-5A bound to the ribosome. Nucleic Acids Res 44:1944–1951. doi:10.1093/nar/gkv1517

Shaffer AL, Shapiro-Shelef M, Iwakoshi NN, Lee A-H, Qian S-B, Zhao H, Yu X, Yang L, Tan BK, Rosenwald A, Hurt EM, Petroulakis E, Sonenberg N, Yewdell JW, Calame K, Glimcher LH, Staudt LM. 2004. XBP1, Downstream of Blimp-1, Expands the Secretory Apparatus and Other Organelles, and Increases Protein Synthesis in Plasma Cell Differentiation. Immunity 21:81–93. doi:10.1016/j.immuni.2004.06.010

Shao S, Murray J, Brown A, Taunton J, Ramakrishnan V, Hegde RS. 2016. Decoding Mammalian Ribosome-mRNA States by Translational GTPase Complexes. Cell 167:1229–1240.e15. doi:10.1016/j.cell.2016.10.046

Sohmen D, Chiba S, Shimokawa-Chiba N, Innis CA, Berninghausen O, Beckmann R, Ito K, Wilson DN. 2015. Structure of the Bacillus subtilis 70S ribosome reveals the basis for species-specific stalling. Nat Commun 6:6941. doi:10.1038/ncomms7941

Sriburi R, Jackowski S, Mori K, Brewer JW. 2004. XBP1. J Cell Biol 167:35–41. doi:10.1083/jcb.200406136

Su T, Cheng J, Sohmen D, Hedman R, Berninghausen O, von Heijne G, Wilson DN, Beckmann R. 2017. The force-sensing peptide VemP employs extreme compaction and secondary structure formation to induce ribosomal stalling. Elife 6:1–17. doi:10.7554/eLife.25642

Svidritskiy E, Brilot AF, Koh CS, Grigorieff N, Korostelev AA. 2014. Structures of yeast 80S ribosome-tRNA complexes in the rotated and nonrotated conformations. Structure 22:1210–1218. doi:10.1016/j.str.2014.06.003

Voorhees RM, Hegde RS. 2015. Structures of the scanning and engaged states of the mammalian SRP-ribosome complex. Elife 4:1–21. doi:10.7554/eLife.07975

Walter P, Blobel G. 1983. Disassembly and reconstitution of signal recognition particle. Cell 34:525–533. doi:10.1016/0092-8674(83)90385-9

Walter P, Ron D. 2011. The Unfolded Protein Response: From Stress Pathway to Homeostatic Regulation. Science 334:1081–1086. doi:10.1126/science.1209038

Wilson DN. 2009. The A-Z of bacterial translation inhibitors. Crit Rev Biochem Mol Biol 44:393–433. doi:10.3109/10409230903307311

Yanagitani K, Imagawa Y, Iwawaki T, Hosoda A, Saito M, Kimata Y, Kohno K. 2009. Cotranslational Targeting of XBP1 Protein to the Membrane Promotes Cytoplasmic Splicing of Its Own mRNA. Mol Cell 34:191–200. doi:10.1016/j.molcel.2009.02.033

Yanagitani K, Kimata Y, Kadokura H, Kohno K. 2011. Translational pausing ensures membrane targeting and cytoplasmic splicing of XBP1u mRNA. Science 331:586–589. doi:10.1126/science.1197142

Yoshida H, Matsui T, Yamamoto A, Okada T, Mori K. 2001. XBP1 mRNA Is Induced by ATF6 and Spliced by IRE1 in Response to ER Stress to Produce a Highly Active Transcription Factor. Cell 107:881–891.

Zhang J, Pan X, Yan K, Sun S, Gao N, Sui SF. 2015. Mechanisms of ribosome stalling by SecM at multiple elongation steps. Elife 4:1–25. doi:10.7554/eLife.09684.001

